# The *C. elegans* 3’-UTRome V2: an updated genomic resource to study 3’-UTR biology

**DOI:** 10.1101/704098

**Authors:** HS Steber, C Gallante, S O’Brien, P.-L Chiu, M Mangone

## Abstract

3’-Untranslated Regions (3’-UTRs) of mRNAs emerged as central regulators of cellular function as they contain important but poorly-characterized *cis*-regulatory elements targeted by a multitude of regulatory factors. The model nematode *C. elegans* is ideal to study these interactions since it possesses a well-defined 3’-UTRome. In order to improve its annotation, we have used a genomics approach to download raw transcriptome data for 1,088 transcriptome datasets corresponding to the entire collection of *C. elegans* trancriptomes from 2015 to 2018 from the Sequence Read Archive at the NCBI. We then extracted and mapped high-quality 3’-UTR data at ultra-deep coverage. Here we describe and release to the community the updated version of the worm 3’-UTRome, which we named 3’-UTRome v2. This resource contains high-quality 3’-UTR data mapped at single base ultra-resolution for 23,084 3’-UTR isoform variants corresponding to 14,788 protein-coding genes and is updated to the latest release of WormBase. We used this dataset to study and probe principles of mRNA cleavage and polyadenylation in *C. elegans*. The worm 3’-UTRome v2 represents the most comprehensive and high-resolution 3’-UTR dataset available in *C. elegans* and provides a novel resource to investigate the mRNA cleavage and polyadenylation reaction, 3’-UTR biology and miRNA targeting in a living organism.

## INTRODUCTION

3’-Untranslated Regions (3’-UTRs) are the portions of mRNA located between the end of the coding sequence and the poly(A) tail of RNA polymerase II-transcribed genes. They contain *cis*-regulatory elements targeted by miRNAs and RNA binding proteins and modulate mRNA stability, localization, and overall translational efficiency (Bartel 2018). Because multiple 3’-UTR isoforms of a particular mRNA can exist, differential regulation of 3’-UTRs has been implicated in numerous diseases, and its discriminative processing influences development and metabolism (Mayr and Bartel 2009; Zhu et al. 2018). 3’-UTRs are processed to full maturity through cleavage of the nascent mRNA and subsequent poly(A) tail addition to its 3’-end by the nuclear poly(A) polymerase enzyme (PABPN1) (Kuhn and Wahle 2004). The mRNA cleavage step is a dynamic regulatory process directly involved in the control of gene expression in eukaryotes. The reaction depends on the presence of a series of sequence elements located within the end of the 3’-UTRs. The most well-characterized sequence is the Poly(A) Signal (PAS) element, a hexameric motif located at ∼19 nt from the polyadenylation site in the 3’-UTR of mature mRNAs. In metazoans, the PAS element is commonly ‘AAUAAA’, which accounts for more than half of all 3’-end processing in eukaryotes (Mangone et al. 2010; Tian and Graber 2012) although alternative forms of the PAS elements exist (Mangone et al. 2010; Jan et al. 2011; Blazie et al. 2015). Previous studies have shown that single base substitutions in this sequence reduce the effectiveness of the cleavage and the polyadenylation of the mRNA transcript (Sheets et al. 1990; Chen et al. 1995). However, this canonical sequence is necessary and sufficient for efficient 3’-end polyadenylation *in vitro* (Clerici et al. 2018; Sun et al. 2018). A less defined ‘GU rich’ element is also known to be present downstream of the cleavage site to facilitate the cleavage and polyadenylation steps (Chen et al. 1995). Recently, studies in human cells identified an additional upstream ‘UGUA’ sequence that is not always present and not required for the cleavage process, but can act as a cleavage enhancer in the context of alternative polyadenylation (APA) if present (Zhu et al. 2018).

APA is a poorly understood mRNA maturation step that produces mRNAs with different 3’-UTR lengths due to the presence of multiple PAS elements within the same 3’-UTR. The usage of the most upstream element, termed the proximal PAS element, leads to the formation of shorter 3’-UTR isoforms while the use of the distal PAS element results in a longer isoform. Importantly, these changes in size may include or exclude regions to which regulatory molecules such as microRNAs (miRNAs) and RNA-binding proteins (RBPs) can bind, substantially impacting gene expression (Matlin et al. 2005; Bartel 2009). While its function in eukaryotes is still not fully understood, a recent study revealed that APA may occur in a tissue-specific manner and, at least in the nematode *C. elegans*, is used in specific cellular contexts to evade miRNA-based regulatory networks in a tissue-specific manner (Blazie et al. 2015; Blazie et al. 2017).

The length of 3’-UTRs is defined during the cleavage and polyadenylation reaction, which is still poorly characterized in metazoans. Although it involves a multitude of proteins and is considered to be very dynamic, the role of each member of the complex and the order in which this process is executed is still not fully understood.

In humans, the Cleavage and Polyadenylation Complex (CPC) is composed of at least 17 proteins (**Figure 1A**) which immunoprecipitate into at least four large sub-complexes: the Cleavage and Polyadenylation Specificity Factor (CPSF), the Cleavage Stimulation Factor (CstF), the Cleavage Factor Im (CFIm) and the Cleavage Factor IIm (CFIIm) sub-complexes (**Figure 1A**). The CPSF sub-complex forms the minimal core component necessary and sufficient to recognize and bind the PAS element of the nascent mRNA *in vitro* (Tian and Manley 2017) (**Figure 1A**). In humans, the CPSF sub-complex is composed of the proteins CPSF160 (Clerici et al. 2017; Sun et al. 2018), CPSF100 (Mandel et al. 2006), CPSF73 (Mandel et al. 2006), CPSF30 (Clerici et al. 2017; Sun et al. 2018), Fip1 (Kaufmann et al. 2004) and Wdr33 (Clerici et al. 2017; Sun et al. 2018). Initial experiments assigned CPSF160 with the role of binding the PAS element, but it is now clear that Wdr33 and CPSF30 are the proteins that instead contact the PAS directly. CPSF160 has a scaffolding role in this process and keeps this sub-complex structured (Chan et al. 2014). The interaction between members of the CPSF core complex (Wdr33, CPSF30, and CPSF160) and the PAS element was recently revealed using single-particle cryo-EM (Clerici et al. 2017; Sun et al. 2018), showing a unique conformation where the PAS element twists to form an s-shaped structure with a non-canonical pairing between the U3 and the A6 in the PAS element (Sun et al. 2018).

**Figure 1.**
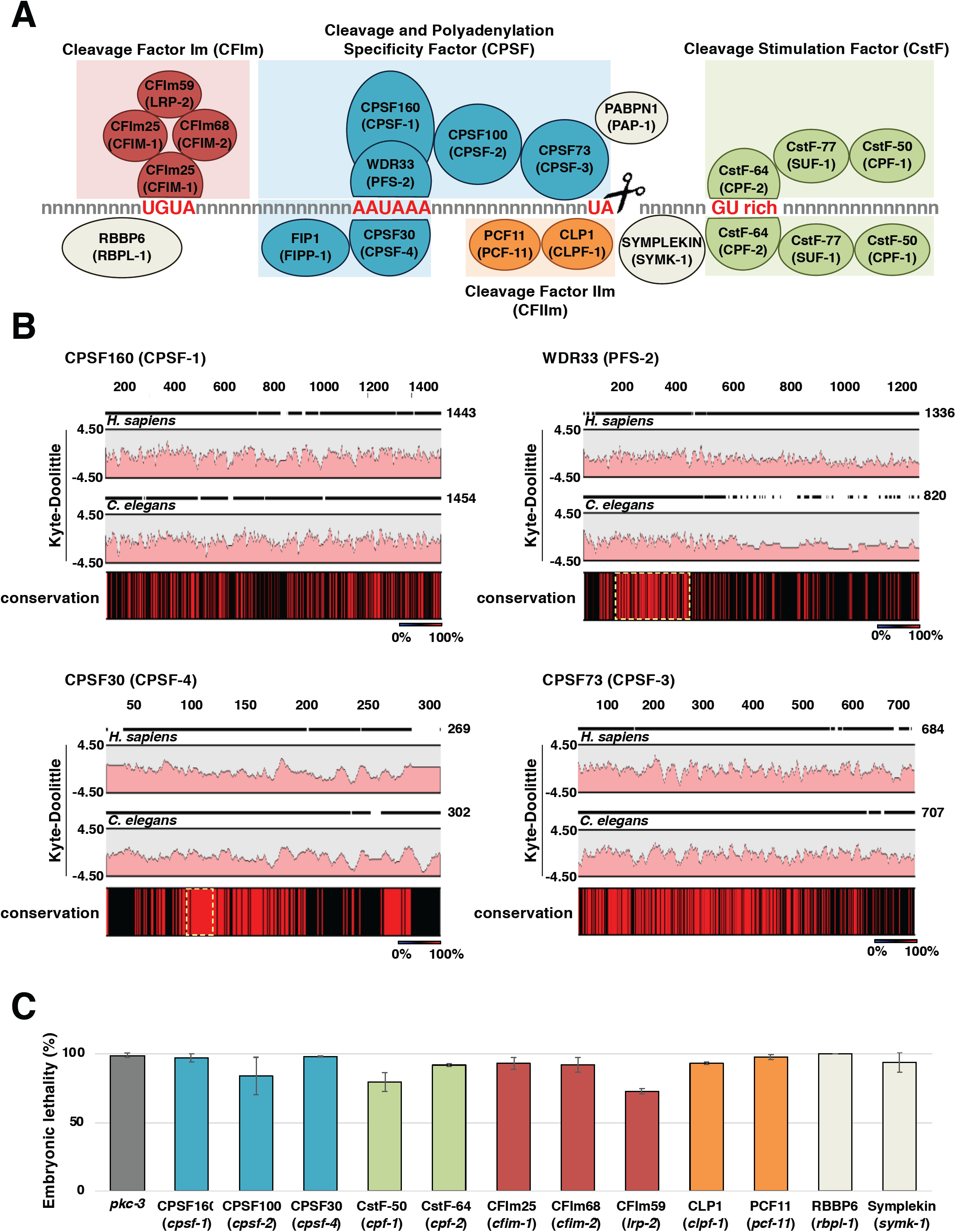
The *C. elegans* members of the Cleavage and Polyadenylation Complex (CPC). A) The CPC is composed of at least 4 independent subcomplexes named Cleavage and Polyadenylation Specificity Complex (Blue), which canonically recognizes the PAS hexamer ‘AAUAAA’ and performs the cleavage downstream of the dinucleotide TA; the Cleavage Stimulation Factor Complex (Green), which binds downstream of the cleavage site to GU rich elements; and the Cleavage Factor CFIm (Red) and CFIIm (Orange) Complexes. CFIm recognizes the element ‘UGUA’ located upstream of the PAS element. This element is not always present. Other known required factors are the Poly(A) Polymerase enzyme, the scaffolding member symplekin and RBBP6. The name of the *C. elegans* orthologs are shown in parenthesis. B) The human and *C. elegans* CPSF subcomplexes are similar in amino acid composition and structure. 2-species alignments between several members of the human and *C. elegans* CPSF members. Amino acids 100% conserved between these two species are shown in red in the conservation bar. Yellow dotted boxes show the sequence of the proteins that interact with the PAS element. Functional domains are conserved. The two Kyte-Doolittle graphs in each panel indicate the hydrophobic amino acids in human and *C. elegans*. C) We have used RNAi to selectively silence most of the members of the CPC complex in *C. elegans*. We observed a strong embryonic lethality phenotype with all the RNAi experiments performed.

CPSF73 is the endonuclease that performs the cleavage of the nascent mRNAs (Ryan et al. 2004; Mandel et al. 2006) (**Figure 1A**). Importantly, CPSF73 is also required in the cleavage of pre-histone mRNAs and is recruited on their cleavage site by the U7 snRNP (Yang et al. 2009).

The CstF sub-complex is the second most well-characterized sub-complex involved in the cleavage and polyadenylation reaction (**Figure 1A**). CstF binds to GU rich elements located downstream of the cleavage site in the nascent mRNA and directly contacts the CPSF sub-complex using its conserved HAT-C domain (Bai et al. 2007; Yang et al. 2018) (**Figure 1A**). The CstF sub-complex is a trimer of heterodimers composed of CstF77, CstF64 and CstF50 (Yang et al. 2018). CstF77 holds the complex together through its Pro-rich domain located on its C terminal region (Takagaki and Manley 2000) **(Figure 1A)**. CstF64 recognizes GU rich sequences through its N-terminal RRM domain (Perez Canadillas and Varani 2003; Yang et al. 2018) and interacts with CstF77 and the scaffolding protein symplekin using its N-terminal HINGE domain (**Figure 1A**) (Takagaki and Manley 2000).

The CFIm and CFIIm sub-complexes are unfortunately less characterized (**Figure 1A**). The CFIm sub-complex is composed of the CFIm68, CFIm59 and CFIm25 subunits, and it was recently shown to contribute to APA by influencing PAS selection (Martin et al. 2012; Hwang et al. 2016). CFIm25 binds the ‘UGUA’ RNA element upstream of the cleavage site and contributes to 3’-end processing and APA by recruiting CFIm59 and CFIm68 (Yang et al. 2010; Yang et al. 2011; Zhu et al. 2018).

Despite the importance of this complex, the CPC remains poorly characterized in most species, including humans, and most of the research in this field is performed *in vitro*.

The roundworm nematode *C. elegans* represents an attractive, novel system to study the cleavage and polyadenylation process *in vivo*. Most of the CPC is conserved between humans and nematodes, including known functional domains and protein interactions (**Figure 1B and Supplemental Figure S1**). *C. elegans* possess the most well-annotated 3’-UTRome available so far in metazoans, with ∼26,000 mapped 3’-UTR boundaries corresponding to ∼16,000 distinct *C. elegans* protein-coding genes (Mangone et al. 2010; Jan et al. 2011).

The *C. elegans* 3’-UTRome was originally developed in 2011 within the modENCODE Project (Mangone et al. 2008; Gerstein et al. 2010; Mangone et al. 2010) and represented a milestone in 3’-UTR biology since it allowed the community to study and identify important regulatory elements such as miRNA and RBP targets with great precision. A second 3’-UTRome was later published using a different mapping pipeline (Jan et al. 2011), confirming most of the previous data such as isoform numbers, PAS usage, etc. Other datasets were made available later, mostly focusing on tissue-specific 3’-UTRs and alternative polyadenylation (Haenni et al. 2012; Blazie et al. 2015; Blazie et al. 2017; Chen et al. 2017; Diag et al. 2018; West et al. 2018).

Although refined and based on several available datasets, only a subset of *C. elegans* 3’-UTRs in protein-coding genes are sufficiently annotated today, and the existing mapping tools do not yet reach the single-base resolution necessary to execute downstream analysis and study the cleavage and polyadenylation process in detail. Most of these 3’-UTR datasets were developed using a gene prediction set now considered obsolete (WS190), and the 3’-UTR coordinates often do not match the new gene coordinates.

To address these and other issues, we developed a novel pipeline to bioinformatically extract 3’-UTR data from almost all *C. elegans* transcriptome datasets stored in the public repository SRA trace archive. This blind approach produced a new saturated dataset we named 3’-UTRome v2, which is available to the community as an additional gBrowse track in the *C. elegans* database WormBase (www.WormBase.org) (Stein et al. 2001; Lee et al. 2018) and in the 3’-UTR-centric database 3’-UTRome (www.UTRome.org) (Mangone et al. 2008; Mangone et al. 2010). We have also used this dataset to study the PAS sequence requirement and the cleavage location of the CPC *in vivo* using transgenic *C. elegans* animals.

## RESULTS

### Functional elements of the human cleavage and polyadenylation complex are conserved in nematodes

To initially gain structural and functional information for the *C. elegans* CPC, we downloaded the protein sequences of the orthologs of the *C. elegans* CPC and aligned them to their human counterparts (**Figure 1B and Supplemental Figure S1**). Based on sequence similarity, *C. elegans* possess orthologs to all the known members of the human CPC, with many peaks of conservation interspersed within known interaction domains of the subunits. The amino acids that make direct contact with PAS elements are also conserved in *C. elegans*; 11 out of the 12 amino acids that form hydrogen bonds and salt bridges with the PAS element (Clerici et al. 2017) are present in both the CPSF30 and WDR33 worm orthologs CPSF-4 and PFS-2 (V67^CPSF30^ with V81^CPSF-4^; K69^CPSF30^ with K83^CPSF-4^; R73^CPSF30^ with R87^CPSF-4^; E95^CPSF30^ with E109^CPSF-4^; K77^CPSF30^ with K91^CPSF-4^; S106^CPSF30^ with S120^CPSF-4^; N107^CPSF30^ with N121^CPSF-4^; R54^WDR33^ with R80^PFS-2^; R47^WDR33^ with R71^PFS-2^; R49^WDR33^ with R73^PFS-2^) (**Figure 1B and Supplemental Figure S1**). The only exception is Y97^CPSF30^, which is substituted with a phenylalanine residue in the worm ortholog. In addition, 9 out of the 10 amino acids in CPSF30 and WDR33 that form the π-π stacking and hydrophobic interactions with the AAUAAA RNA element (Clerici et al. 2017) are also conserved in the CPSF30 and WDR33 worm orthologs CPSF-4 and PFS-2 (A1:K69^CPSF30^ with K83^CPSF-4^ and F84^CPSF30^ with F98^CPSF-4^; A2: H70^CPSF30^ with H84^CPSF-4^; U3: I156^WDR33^ with I181^PFS-2^; A4: F112^CPSF30^ with F126^CPSF-4^ and F98^CPSF30^ with F112^CPSF-4^; A5: F98^CPSF30^ with F112^CPSF-4^; A6: F153^WDR33^ with F178^PFS-2^) (**Figure 1B and Supplemental Figure S1**). The only exception is a F43WDR33 substitution to a glycine residue that interacts with A6 in the worm ortholog.

CPSF73, the endonuclease that performs the cleavage reaction, has a *C. elegans* ortholog named CPSF-3. Both genes are conserved with an overall 57.61% identity that increases to 69.52% in the β-lactamase domain, which is the region required to perform the cleavage reaction (**Figure 1B and Supplemental Figure S1**). Specifically, all eight amino acids shown previously to form the zinc binding site required for the cleavage reaction (Mandel et al. 2006) are also conserved (D75^CPSF73^ with D74^CPSF-3^; H76^CPSF73^ in H75^CPSF-3^; H73^CPSF73^ in D72^CPSF-3^; H396^CPSF73^ with H397^CPSF-3^; H158^CPSF73^ with H159^CPSF-3^; D179^CPSF73^ with D180^CPSF-3^; H418^CPSF73^ with H419^CPSF-3^; E204^CPSF73^ with E205CPSF-3) (**Figure 1B and Supplemental Figure S1**). This overall similarity is also observed in most of the other members of the *bona fide C. elegans* CPC complex (**Supplemental Figure S1**), suggesting similar structure and function.

In addition, when subjected to RNAi analysis, each of the *C. elegans* CPC members produced a similar strong embryonic lethal phenotype, suggesting that each of these genes may act as a complex and is required for viability (**Figure 1C and Supplemental Figure S2)**.

### An updated 3’-end mapping strategy

Next, we used a genome-wide approach to improve the current version of the 3’-UTRome. We refined a 3’-UTR mapping pipeline we previously developed and used in the past (Blazie et al. 2015; Blazie et al. 2017). This approach uses raw transcriptome data as input material to identify and precisely map high-quality 3’-UTR end clusters (**Figure 2 and Supplemental Figure S3**). A similar approach has been previously applied to study *C. elegans* transcriptomes in the past (Tourasse et al. 2017).

**Figure 2.**
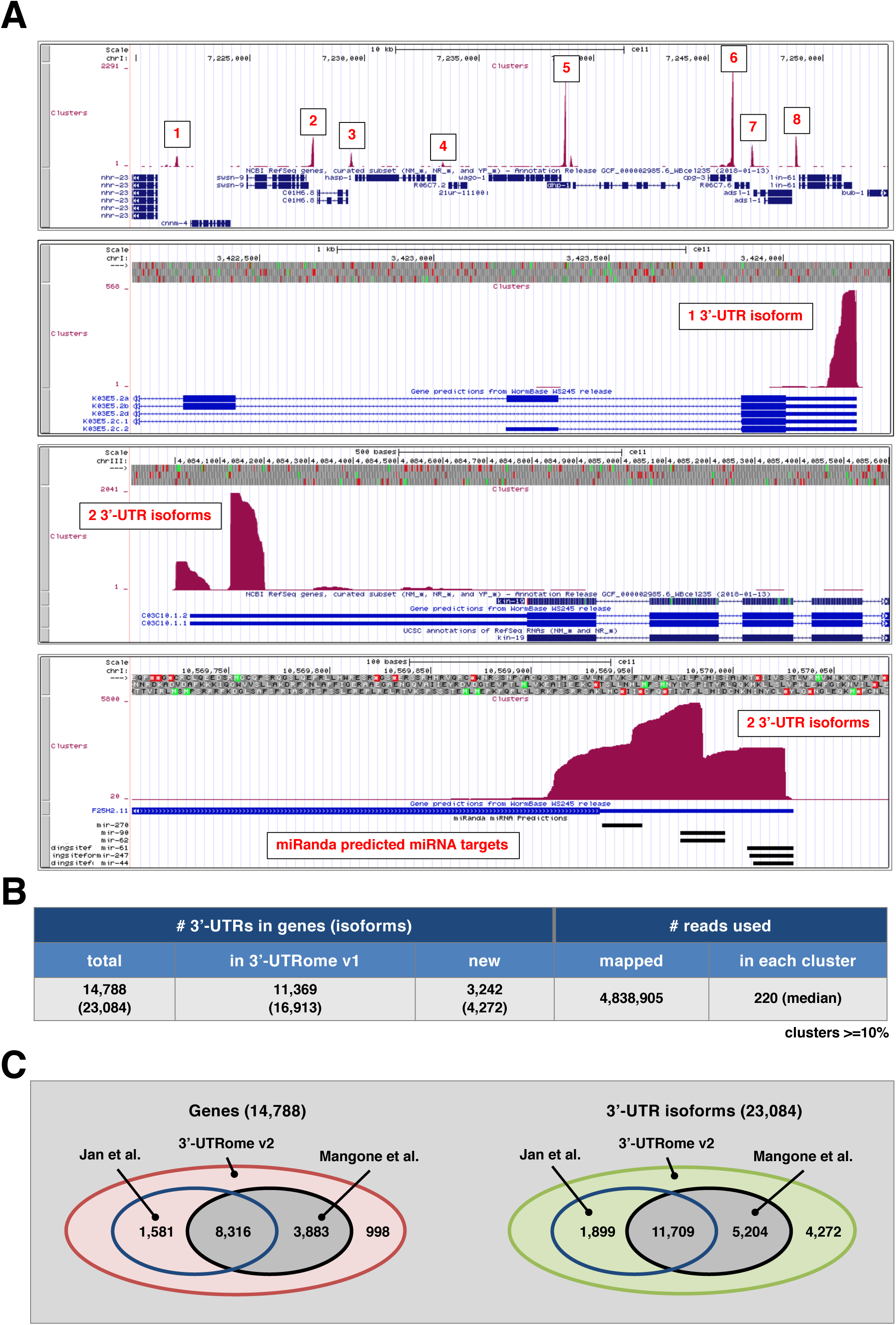
Cluster preparation and analysis. A) Screenshots showing several mapped 3’-UTR clusters for genes with one or two 3’-UTR isoforms. MiRanda predicted miRNA targets are shown for a particular 3’-UTR at the bottom of this Panel. B) Summary of the 3’-UTRs in genes identified in this study along with the number of reads mapped and clustered for each 3’-UTR. C) Comparison between the 3’-UTRs for genes and total isoforms mapped in this study vs the UTRome v1 (Mangone et al. 2010) and the dataset from Jan et al. (2011).

We wanted to obtain the most accurate, saturated and tissue-independent dataset possible. To achieve this goal we downloaded the entire collection from 2015 to 2018 of transcriptome datasets stored in the Sequence Read Archive (SRA) (**Supplemental Table S1**) and processed them through our 3’-UTR mapping pipeline. We reasoned that this approach would lead to the identification of as many 3’-UTR isoforms as possible in an unbiased manner since these downloaded transcriptomes have been sequenced using both *wild-type* and mutant strains subjected to many different environmental conditions and covering all developmental stages with many replicates.

We downloaded a total of 1,088 *C. elegans* transcriptome datasets (∼2TB of total raw data) (**Supplemental Table S1**). Most of these datasets have also been used in the past to map polyadenylation sites in *C. elegans*. Our 3’-UTR mapping approach extracted from these datasets ∼5M unique, high-quality poly(A) reads, which we then used for cluster preparation and mapping (see Methods). We implemented very restrictive parameters for cluster identification and 3’-UTR end mapping to limit the unavoidable noise produced by using such diverse datasets as data sources (**Supplemental Figure S3**). Our approach led us to map 3’-UTR clusters with ultra-deep coverage of several magnitudes (average cluster coverage ∼220X) (**Figure 2A**) and to identify 23,084 3’-UTR isoforms corresponding to 14,788 protein-coding genes. When compared to the previous 3’-UTRome v1 dataset (Mangone et al. 2010), we obtained 3’-UTR information for an additional 3,242 new protein-coding genes (4,272 3’-UTR isoforms) (73% of all protein-coding genes included in the WS250 release) (**Figure 2B-C and Supplementary Figure S4**).

### The *C. elegans* 3’-UTRome v2

Our approach produced high-quality 3’-UTR data for 14,788 *C. elegans* protein-coding genes (**Figure 2B**). The most abundant nucleotide in *C. elegans* 3’-UTRs is a uridine, which accounts for ∼40% of all nucleotides in 3’-UTRs (**Figure 3A Top Left Panel**). Adenosine nucleotides are the second most represented nucleotide class with ∼30% incidence (**Figure 3A Top Left Panel**). Alternative polyadenylation is common but occurs at a lesser extent than what was previously published (Mangone et al. 2010; Jan et al. 2011). The majority of protein-coding genes (58%) are transcribed with only one 3’-UTR isoform (**Figure 3A Bottom Left Panel**) which closely resembles the previously reported ∼61% (Mangone et al. 2010; Jan et al. 2011). Genes with two 3’-UTR isoforms are notably increased in occurrence when compared with past studies (32% vs 25%), while the occurrence of genes with three or more 3’-UTRs is comparable with what was previously found (**Figure 3A Bottom Left Panel**) (Mangone et al. 2010; Jan et al. 2011).

**Figure 3.**
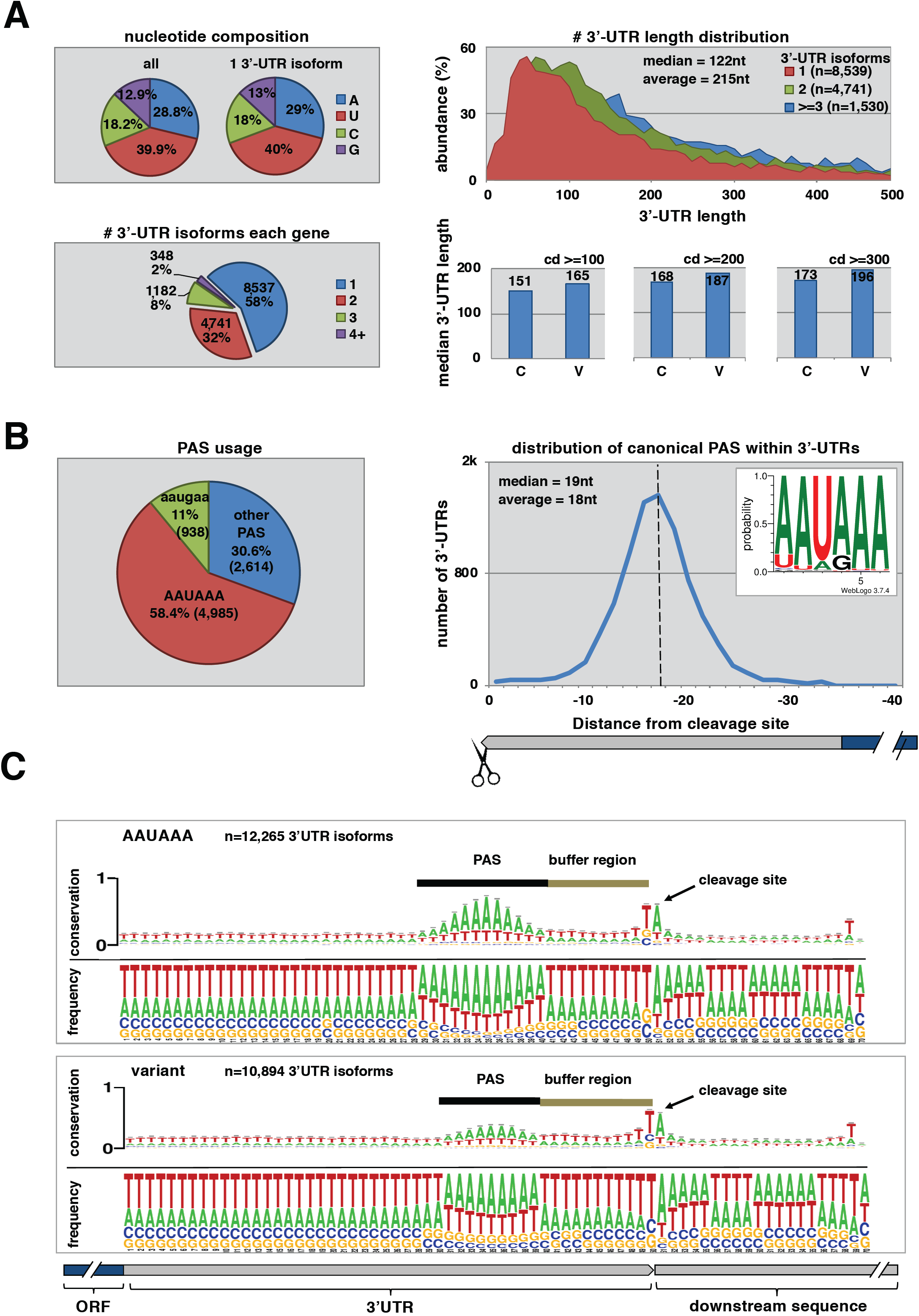
The worm 3’-UTRome v2. A) Top Left Panel. Nucleotide composition of 3’-UTRs in the 3’-UTRome v2. Uridine is the most abundant nucleotide within 3’-UTRs for *C. elegans*. Bottom Left Panel. The number of 3’-UTR isoforms in each gene. 42% of the genes in the 3’-UTRome v2 possess multiple 3’-UTR isoforms. Top Right Panel. 3’-UTR length distribution in genes expressed with one, two, or three or more 3’-UTR isoforms. The median 3’-UTR length across these datasets is 122 nt. Genes with multiple 3’-UTR isoforms are on average longer than genes with one 3’-UTR isoform. Bottom Right Panel. Median 3’-UTR length in genes with Canonical (C) or Variant (V) PAS elements. There is a slight increase in 3’-UTR length in genes with variant PAS elements when compared to those with canonical PAS elements. This variation is still detected when increasing the stringency of the density of the clusters (cd) used in this analysis. B) Left Panel. PAS element usage in 3’-UTRs. 58.4% of 3’-UTRs use the canonical PAS element ‘AAUAAA’ while the most common variant PAS element is the hexamer ‘AAUGAA’, which occurs in 11% of genes. Right Panel. The distribution of canonical PAS elements within 3’-UTRs. The average distance from the PAS element to the cleavage site is 18 nt. C) Alignment of 3’-UTRs at the cleavage site. This alignment in genes with both canonical and variant PAS elements reveals a region between the PAS element and the cleavage site we renamed the *buffer region* in which cleavage rarely occurs. The most abundant nucleotide at the cleavage site is an adenosine nucleotide preceded by a uridine nucleotide.

In the case of genes with multiple 3’-UTRs, the canonical AAUAAA PAS site is over two times more abundant in longer 3’-UTR isoforms than in shorter 3’-UTR isoforms, suggesting that the preparation of shorter 3’-UTR isoforms may be subject to regulation (**Supplemental Figure S5**).

The mean 3’-UTR length in the 3’-UTRome v2 is 215 nt (**Figure 3A Top Right Panel**), and the occurrence of more 3’-UTR isoforms per gene correlates with an overall extension in length (**Figure 3A Top Right Panel**). We also note a slight correlation between 3’-UTR length and PAS element usage, with longer 3’-UTRs more frequently containing variant PAS elements (**Figure 3A Bottom Right Panel**). The most common PAS element in *C. elegans* protein-coding genes is consistently the hexamer ‘AAUAAA’, which is present in 58.4% of all the 3’-UTRs mapped in this study (**Figure 3B Left Panel**). This element is ∼20% more abundant than what was previously identified in past studies (Mangone et al. 2010; Jan et al. 2011), and a slight variation of this motif is also present in genes with no canonical PAS elements (**Supplemental Figure S6**). The PAS sequence is located ∼18 nt from the cleavage site (**Figure 3B Right Panel**), and a *buffer region* of ∼12 nt is present between the PAS element and the cleavage site (**Figure 3C**). The cleavage site occurs almost invariably at an adenosine nucleotide, which is often preceded by a uridine nucleotide (**Figure 3C**). A GO term analysis in genes with 1, 2 or 3 3’-UTR isoforms (**Supplemental Figure S7**) revealed a few unique patterns with no major hits, perhaps because APA is so widespread in *C. elegans* and affects almost half of its known protein coding genes.

### An RRYRRR motif in 3’-UTRs with variant PAS elements

We could not detect any enrichment for the UGUA motif near the cleavage site (**Supplemental Figure S8**). Perhaps this element is not used in *C. elegans*, or the CFIm complex may recognize a variant motif not yet identified in this organism. Importantly, when we aligned the 3’-ends of 3’-UTRs which contain variant PAS elements, we noticed an enrichment of a ‘RRYRRR’ motif which in most instances resembles a canonical AAUAAA motif with a guanine replacing the A4 nucleotide (**Figure 4A and Supplemental Figure S6**). This finding suggests that in *C. elegans* a ‘RRYRRR’ element could be used when the AAUAAA hexamer is absent (**Figure 4A**). We have also identified other conserved elements which need to be further validated (**Supplemental Figure S9**)

**Figure 4.**
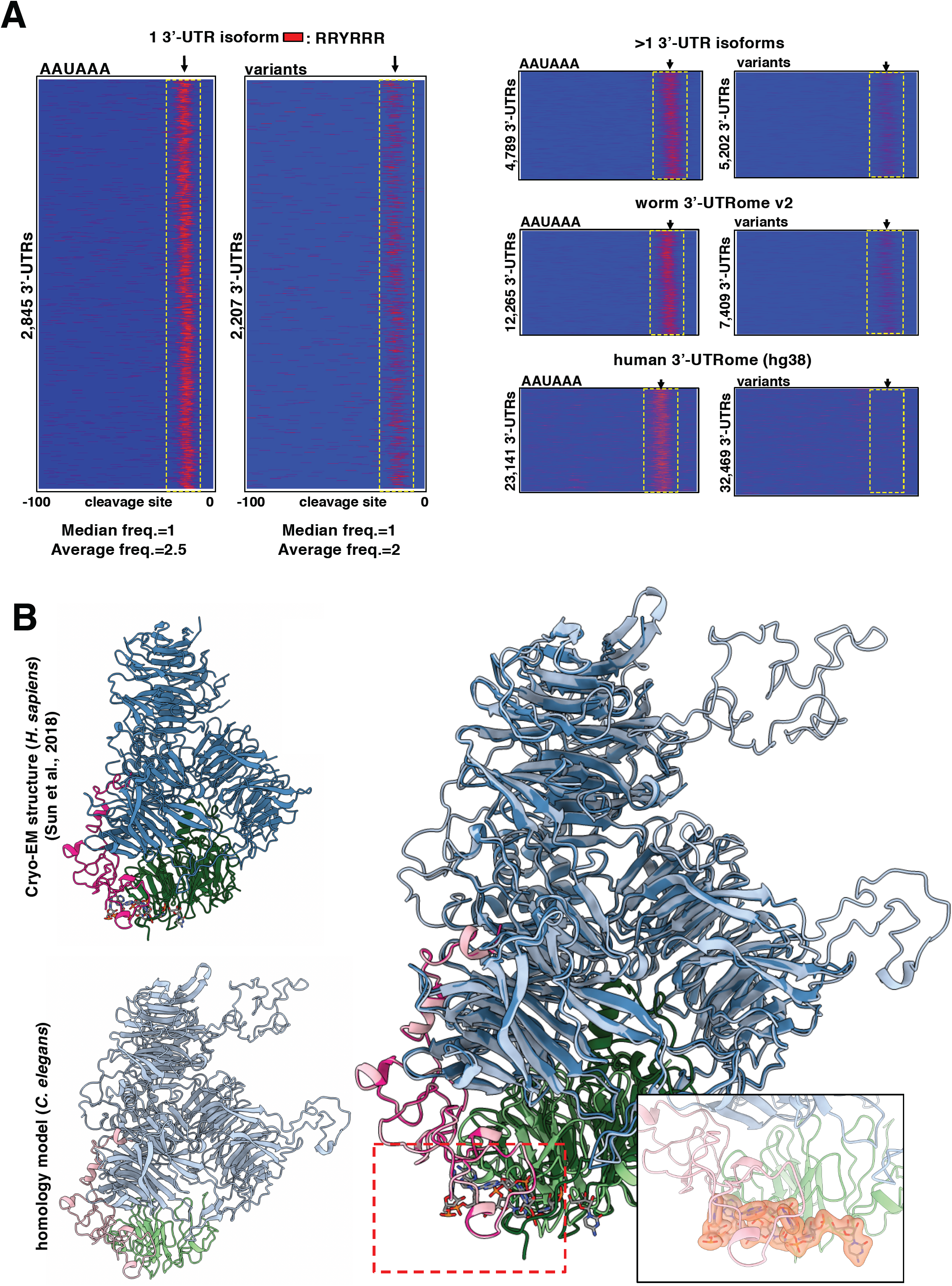
The sequence requirements of the *C. elegans* CPSF core complex. A) PAS element usage of the RRYRRR motif. 3’-UTRs from the 3’-UTRome v2 aligned by their cleavage site in genes with canonical or variant PAS element. Instances of the motif RRYRRR are represented by the thin red bars mapped on 3’-UTR sequences with the coordinates −100 nt to 0 nt, where 0 nt represents the cleavage site. The spatial conservation highlighted by the yellow box of this RRYRRR motif is very strong in single 3’-UTR isoforms with canonical PAS elements and is enriched in those with variant PAS elements. This RRYRRR element is maintained in 3’-UTRs that have at least two isoforms, but it is not strongly represented in human 3’-UTR data (hg38) due to the lack of their annotation. R= purine, Y= pyrimidine. B) Superimposition of the cryo-EM structure of the previously published human CPSF core complex (Clerici et al. 2018; Sun et al. 2018) to the worm CPSF core complex: cpsf-1 (CPSF-160) in blue, pfs-2 (WDR33) in pink, and cpsf-4 (CPSF30) in green. The PAS element binding pocket can be fitted into the homology model. The PAS element of the RNA is represented in yellow. The size and the selectivity of the nucleotide binding pocket can fit other nucleotides as long as the motif is RRYRRR.

To better understand the molecular details of the interaction between CPSF and the PAS element, we built an atomic homology model of the worm CPSF core complex containing CPSF-1 (CPSF160), PFS-2 (Wdr33), and CPSF-4 (CPSF30) (**Figure 4B and Supplemental Figure S10)**. Most of this model can be superimposed to the cryo-EM structure of the human CPSF core complex (**Figure 4B and Supplemental Figure S10**).

The nucleotide-binding pocket can also be fitted into our homology model, which may implicate a conserved binding region in the *C. elegans* complex (**Figure 4B Right Panel)**. From the structural details of the human CPSF core complex, the interactions between the RNA nucleotides and CPSF30 or WDR33 are not specific. The nucleotide binding is mainly established by the π-π ring stacking force between the nucleotide bases and the residues with aromatic side chains, such as phenylalanine and tyrosine (**Supplemental Figure S10**). Also, the buried area of the nucleotide-binding sites in our model was 1138 Å, which is similar to the nucleotide-binding pocket in the human complex (1261 Å). The RMSD of the two models (1.170 Å) indicates a high structural similarity. As observed in the cryo-EM structure by Sun et al. (2018), no specific interactions between nucleotides and the adjacent residues were found, and the interactions between the nucleotide and adjacent residues’ side chains are mostly π-π ring stacking force (**Supplemental Figure S10**). The actual interactions between the bound nucleotides and proteins will need to wait for the structure of the complex determined by crystallography or cryo-EM to validate it. (**Supplemental Figure S10**). Thus, at least in *C. elegans*, the selectivity of the nucleotide binding may be only at a level to the nucleotide bases, that is, pyrimidines or purines.

### An enrichment of adenosine nucleotide at the cleavage site

We were intrigued by the almost invariable presence of adenosine nucleotides near the cleavage site. This enrichment becomes more evident when we sort 3’-UTRs with canonical PAS elements by the length of their respective *buffer regions* (**Figure 5A**). In the case of the largest group with a *buffer region* of 12-13 nt, more than 2,000 3’-UTRs terminate with ∼70% occurrence of adenosine nucleotides at the cleavage site preceded most of the time by a uridine. Since we bioinformatically removed the poly(A) sequences from the sequencing reads during our cluster preparation step, we do not have direct evidence that this last adenosine nucleotide is indeed present in the mature transcripts and used as a template for the polymerization of the poly(A) tail or that it is attached by PABPN1 during the polymerization of the poly(A) tail. Of note, the high abundance of this nucleotide at the cleavage site suggests that it is somehow important in the cleavage process.

**Figure 5.**
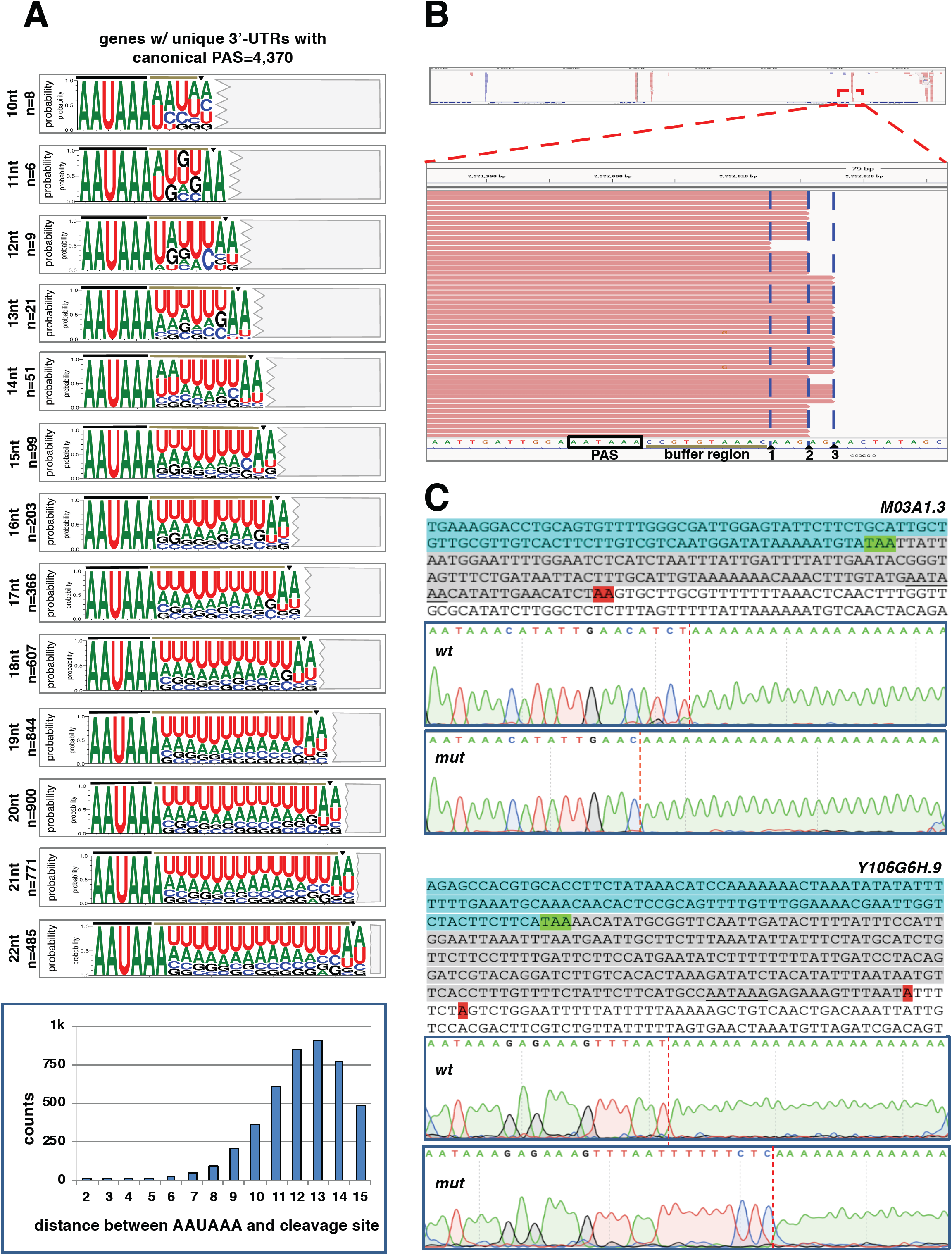
A terminal adenosine nucleotide is required at the cleavage site for correct cleavage. A) Sequence Logos produced from 3’-UTRs of genes only with 3’-UTR isoforms containing the canonical PAS element ‘AAUAAA’ and aligned by their respective *buffer region* length (n=4,374). Two extra nucleotides are included downstream of each cut site (triangle), highlighting the terminal UA dinucleotide. The nucleotide distribution of the distance between the PAS element and the cleavage site is shown in the bar chart below. B) Example of slight variability in the cleavage site for the gene C09G9.8. While prevalent forms are observed, the exact cleavage site varies on several occasions, but it predominantly occurs at a different adenosine nucleotide. C) Test of the role of the terminal adenosine nucleotide in the cleavage reaction. The 3’-end regions of several test genes were cloned and used to prepare transgenic *C. elegans* strains expressing this region with or without mutated terminal adenosine nucleotides (Red, see below). The top sequence shows the test 3’-end region (Cyan=ORF, Green=translation STOP signal, Grey=3’-UTR, Red=Terminal adenosine nucleotide. The PAS element is underscored). The Sanger trace files show the outcome of the cleavage site location in selected clones. Two genes are shown (M03A1.3 and Y106G6H.9). In the case of M03A1.3, the loss of the terminal adenosine nucleotide sometimes forces the CPC to backtrack and cleave the mRNAs upstream of the regular cleavage site but still at the closest adenosine nucleotide available. In the case of the gene Y106G6H.9, the loss of the terminal adenosine nucleotide forces the complex to skip the cleavage site, which sometimes occurs at the next purine nucleotide. Additional clones and more test genes are shown in **Supplemental Figures S11-S13**.

We decided to investigate this issue further and study how precisely the raw reads produced by our cluster algorithm align to the genome. We noticed that in each gene, the cleavage rarely occurs at a unique position in the transcript. Instead, there are always slight fluctuations of the exact cleavage site, with a few percentages of reads ending a few nucleotides upstream or downstream of the most abundant cleavage site for a given gene (**Figure 5B**). Importantly, almost all the reads in each cluster terminate at an adenosine nucleotide (**Figure 5B**). Also, if there are adenosine nucleotides located within shorter *buffer regions*, the cleavage rarely occurs at these sites. Perhaps, the large size of the CPC does not allow for the docking and the cleavage of the pre mRNAs near the PAS element, which is optimally performed at 12-13 nt downstream of the PAS element (**Figure 5A and Figure 5B**).

Next, we decided to study the role of the terminal adenosine nucleotide in the cleavage process. We reasoned that if this adenosine nucleotide indeed plays any role in the cleavage process, we should be able to alter the position of the mRNA cleavage site by mutating this residue with different purines or pyrimidines in the pre mRNAs of selected test genes.

We selected three test genes; *ges-1*, Y106G6H.9, and M03A1.3. These genes are processed with a single 3’-UTR isoform, use a single canonical PAS element, have a *buffer region* of 12, 13 and 14 nucleotides respectively, and possess a terminal uridine and adenosine nucleotide in their sequence. To capture their entire 3’-UTR region, we cloned the genomic portions of these genes spanning from their translation STOP codons to ∼200 nt downstream of their cleavage sites. We then prepared several mutant *C. elegans* strains by replacing their terminal adenosine nucleotide at their cleavage site with other nucleotides. In the case of Y106G6H.9, we also prepared a double mutant removing an additional adenosine nucleotide downstream of the first one located at the cleavage site (**Figure 5C and Supplemental Figure S11-S13**).

We cloned these *wt* and mutant 3’-UTR regions downstream of a GFP reporter vector and prepared transgenic *C. elegans* strains that express them in the worm pharynx using the *myo-2* promoter. We opted to use the pharynx promoter since it is very strong and produces a robust expression of our constructs (**Supplemental Figure S11-S13**). We prepared transgenic worm strains expressing these constructs, recovered total RNAs, and tested if the absence of the terminal adenosine nucleotide in our mutants affects the position of the cleavage site using RT-PCR and a sequencing approach (**Figure 5C and Supplemental Figure S11-S13**).

We observed an overall disruption of the cleavage process, in some case more pronounced than in others (**Figure 5C and Supplemental Figure S11-S13**). In the case of M03A1.3, the absence of the terminal adenosine nucleotide forces the cleavage complex to backtrack in 40% of the tested clones and cleave the mRNAs 3 nt upstream of the original cleavage site but still at an adenosine nucleotide (**Figure 5C and Supplemental Figure S11**). Of note, the new cleavage site does not possess the conserved uridine upstream of the adenosine residue, suggesting that perhaps this nucleotide is not strictly required for the cleavage reaction.

In the case of Y106G6H.9, the single mutant does not alter the position of the cleavage site, but it activates a novel cryptic cleavage site 100 nucleotides upstream of the canonical cleavage site in 20% of the sequenced clones (**Figure 5C and Supplemental Figure S12**). This new site also possesses a terminal UA dinucleotide, a non-used PAS element containing the motif YRYRRR, which could still be recognized by the CPSF core complex, and a *buffer region* of 12 nt. In one case, the Y106G6H.9 double mutant skips the original cleavage site but still cut at the next purine residue, which is not an adenosine in this case (**Supplemental Figure S12**). In the case of *ges-1*, mutating the terminal adenosine does not change the cleavage pattern, although it became more imprecise (**Supplemental Figure S13**).

### Updated miRanda prediction in *C. elegans*

Next, we used our new 3’-UTRome v2 dataset to update miRanda miRNA target predictions. We downloaded and locally ran the miRanda prediction software (John et al. 2004) using our new 3’-UTRome v2 as a target dataset. We have produced two sets of predictions: one generic, which contains the entire output produced by the software, and one more restrictive, which contains only output predictions with high scores and with low E-energy scores. These two tracks have been uploaded in both the 3’-UTRome database (Mangone et al. 2008; Mangone et al. 2010) and WormBase (Lee et al. 2018). Alternative polyadenylation has been previously shown to allow genes to evade miRNA regulation (Blazie et al. 2017). To study this process in the context of miRNA targeting, we have also performed a GO term analysis on the genes known to use APA that either lose or gain a miRNA target (**Supplemental Figure S14**). We have uploaded this dataset as a **Supplementary Table S4**.

## DISCUSSION

Here we have used a genome-wide approach to refine and study the 3’-UTRome in the nematode *C. elegans*. We have identified 3’-UTR data for 14,788 genes corresponding to 23,084 3’-UTR isoforms, improving their annotation. We now have 3’-UTR data for 73% of all protein-coding genes included in the WS250 release. This dataset is not complete, since we could not assign 3’-UTR data for the remaining 5,554 protein-coding genes present in WS250 (**Supplemental Table S4**). For the majority of these genes, their 3’-UTR data were discarded by our highly stringent filters used during 3’-UTR cluster preparation. In addition, some of these genes may be transcribed at very low abundance and their mRNA is present below the sensitivity of our approach. Further experiments need to be performed to identify 3’-UTR data for the remaining 5,554 protein-coding genes.

Transcriptome data does not always reach the resolution needed to map 3’-ends of mRNAs at single base resolution, since reads containing poly(A)s close to the cleavage site are generally discarded by aligners. In the case of short 3’-UTRs which overlap entirely with a single sequencing read, it is possible to successfully attach a given 3’-UTR cluster to the correct gene. However, since the majority of 3’-UTRs in *C. elegans* are longer than the average length of a single read, we do not have a continuous coverage from the translation STOP site to their 3’-end for most of our 3’-UTRs. To attach our clusters to a given gene we rely on a common practice which bioinformatically attaches them to the closest gene within 2,000 nt in the correct orientation (Mangone et al. 2010; Jan et al. 2011).

Alternative polyadenylation is widespread in *C. elegans*, with ∼42% of genes possessing at least two 3’-UTR isoforms (**Figure 3A**). The PAS usage is still most commonly the hexamer ‘AAUAAA’ which is used to process ∼58% of all *C. elegans* 3’-UTRs (**Figure 3B**). Importantly, we found that the remaining 42% possess a variation of this canonical PAS element which indeed is very similar in chemical composition and contains an ‘RRYRRR’ motif at the same location where the PAS element is expected (**Figure 4A**). We do not have direct evidence that the CPC recognizes this motif, but, since it is so conserved, we hypothesize that in *C. elegans* it may provide a docking site in the absence of the canonical AAUAAA site during the cleavage reaction. Additional elements in the buffer region may play a role in this process, but unfortunately this region is very rich in Uridine residues (**Figure 3C** and **Figure 5A**), which makes the identification of the conserved signatures using common motif search software (Bailey et al. 2009) challenging. Importantly, when we studied the sequence requirement of this RRYRRR motif in 3’-UTRs of genes without a canonical PAS element (**Supplemental Figure S6**), we found that the most conserved nucleotides are the Y3 and R6. These two nucleotides are adjacent to each other when bound to WDR33 and CPSF30 in human (Sun et al. 2018) and form a Hoogsteen U-A base pair. This interaction is perhaps required to lock the mRNA in place by these two factors and is size-dependent.

Our superimposition of the *C. elegans* CPSF ortholog to the human cryo-EM structure (Clerici et al. 2018; Sun et al. 2018) in **Figure 4B and Supplemental Figure S10**, although not reinforced by experimental data, still supports our hypothesis, suggesting that in worms the pocket used by this complex to bind the PAS element may accommodate other nucleotides as long as they have a similar chemical structure and can recapitulate the ‘RRYRRR’ motif. In humans, the second most abundant PAS element is ‘AUUAAA’ (Sun et al. 2018), which does not follow this guideline, suggesting that perhaps other factors can contribute to the cleavage of non-canonical PAS elements in other species.

Our analysis on the cleavage site found that the cleavage and polyadenylation machinery does not always cleave the same mRNA at the same position on the 3’-UTR (**Figure 5B**). While a predominant site is often chosen for each gene, a slight variation of a few nucleotides upstream or downstream of the cleavage site is also possible. Importantly, this slight variation almost invariably ends at an adenosine nucleotide in the genome, suggesting that this nucleotide is somehow ‘sensed’ in the cleavage process.

Our mutagenesis results also support an important role for the terminal adenosine nucleotide during the cleavage reaction (**Supplemental Figures S11-S13**). In those experiments, the loss of this terminal adenosine nucleotide disrupts the location of the cleavage in some cases, either activating cryptic cleavage sites or backtracking and using a different adenosine nucleotide upstream of the canonical cleavage site (**Supplemental Figures S11-S13**). Unfortunately, we did not mutate the upstream uridine residue, and we do not know its contribution, if any, to the cleavage reaction. Although we always detected its presence at the cleavage site (except in one case), more experiments need to be performed to confidently assign a role in this process.

The concept of mRNAs terminating with an adenosine nucleotide is not novel. Pioneering work using 269 vertebrate cDNA sequences has shown that ∼71% of these genes terminate with a CA dinucleotide element (Sheets et al. 1990). These experiments were biochemically validated a few years later using SV40 Late Poly(A) signal in mammalian cells in a more controlled environment (Chen et al. 1995). These experiments also showed that, at least for the case of this specific 3’-UTR, the cleavage could not occur closer than 11 nt or further than 23 nt from the PAS element (Chen et al. 1995). In this context, these findings could explain why we do not detect a terminal adenosine at the cleavage site with our double mutant Y106G6H.9, which is 27 nt downstream of the PAS element (**Supplemental Figure S12**). In the case of this gene, the cleavage still occurs at a purine nucleotide, which suggests that perhaps another terminal purine can compensate for the absence of an adenosine nucleotide.

Overall, experiments in **Figure 5C and Supplemental Figures S11-S13** support and expand both these initial results, showing that altering the nucleotide composition downstream of the PAS element may influence the location of the cleavage.

Unfortunately, our study does not have the resolution to definitely verify if this adenosine nucleotide is indeed included in the processed mRNAs or used by the CPC as a genomic mark of the cleavage site. More specifically we do not know if this nucleotide is read by the RNA polymerase II and incorporated in the nascent mRNAs or if the machinery somehow ‘senses’ its presence and cleaves the mRNA upstream of it. Another plausible hypothesis is that CPSF73 may cleave the mRNAs somewhere downstream of this terminal adenosine nucleotide, and then unknown exonucleases degrade the mRNA molecule until the first adenosine in a row is reached. Some insights may come from the process underlining histone 3’-end formation, since CPSF73 also cleaves these poly(A)-lacking histone mRNAs. In this specific case, the enzyme is positioned near the cut site by the U7 snRNP and cuts the nascent pre-mRNA just downstream of an adenosine nucleotide (Yang et al. 2009). We speculate that perhaps CPSF73 is capable of either ‘sensing’ this terminal adenosine nucleotide or is positioned next to it by either other members of the CPC or a not yet identified factor.

If this terminal adenosine is indeed incorporated in the pre-mRNAs, its functional requirement is unclear. It may be used by the poly(A) polymerase enzyme as a substrate to extend the poly(A) tail after the cleavage reaction has been completed or perhaps has an unknown regulatory function. More experiments need to be performed to answer these questions.

While we observed a terminal adenosine nucleotide in most of the mapped 3’-UTRs, the cytosine nucleotide previously identified upstream of the terminal adenosine in humans is replaced with another pyrimidine nucleotide in *C. elegans* (uridine) (**Figure 3C**), suggesting that other factors may contribute to the cleavage site decision by the CPC in higher eukaryotes.

MiRanda predictions were obsolete and needed to be updated since those present in the microrna.org database (www.microrna.org) were obtained using a 9-year-old 3’-UTR dataset. Also, before this study WormBase (Lee et al. 2018) did not include miRNA targeting predictions in its JBrowse software.

The number of predicted miRNA targets is now decreased from 34,186 to 23,160, mostly because several 3’-UTR isoforms in the 3’-UTRome v1 were discarded in this new 3’-UTRome v2 release. We used these new predictions to detect several instances of genes which use APA and can potentially escape miRNA targeting (**Supplemental Table S3**).

In conclusion, this new 3’-UTR dataset, which we renamed 3’-UTRome v2 (**Supplemental Table S5**), has been uploaded to WormBase WS274 release (Lee et al. 2018) and is shown as a new track in the JBrowse tool together with updated miRanda miRNA targets. The 3’-UTRome v2 expands the old 3’-UTRome developed within the modENCODE Consortium, and, together with updated miRanda predictions, provides the *C. elegans* community with an important novel resource to investigate the RNA cleavage and polyadenylation reaction, 3’-UTR biology and miRNA targeting.

## METHODS

### 3’-UTR mapping pipeline

We have used the SRA toolkit from the NCBI to download raw reads from 1,094 transcriptome experiments. The complete list of datasets used in this study is shown in **Supplemental Table S1**. We restricted the analysis to sequences produced from *C. elegans* transcriptomes using the Illumina platform with reads of at least 100 nt in length. At the completion of the download step, the files were unzipped and stored in our servers. We then used a custom-made Perl script to extract reads containing at least 23 consecutive adenosine nucleotides at their 3’-end or 23 consecutive thymidine nucleotides at their 5’-end (**Supplemental Code**). This filter produced 24,973,286 mappable 3’-end reads. We then removed the terminal adenosine or thymidine nucleotides from these sequences, converted them to FASTQ files using the FASTX-Toolkit (CSHL), and mapped them to the WS250 release of the *C. elegans* genome using Bowtie 2 algorithm with standard parameters (Langmead and Salzberg 2012). Bowtie 2 mapped 7,761,642 reads (31.08%), which were sorted and separated based on their respective strand origin (positive or negative). We have uploaded to WormBase two versions of this dataset. The more stringent one, which we named ‘filtered’, includes all these aforementioned filters and has been used in all the analyses performed in this study. A second dataset, named ‘mild’, includes 3’-UTR isoforms which overlap +/-2 nt and have cluster reads <5. The complete set of 3’-UTRs composing the 3’-UTRome v2 are shown in **Supplemental Table S5**.

### Cluster Preparations

Poly(A) clusters were prepared as follow. We stored the ID, genomic coordinates, and strand orientation of each mapped read and used this information throughout the pipeline. The BAM file produced by the aligners was sorted and converted to BED format using SAMtools software (Li et al. 2009). Contiguous genomic coordinates were merged using BEDTools software (Quinlan and Hall 2010) using the following command `‘Bedtools merge -c 1 -o count -I > tmp.cluster’.` This new file produced the characteristic ‘shark fin’ graph visible in **Figure 2**. We used several stringent filters to eliminate as much noise as possible. 1) We ignored clusters composed of less than 6 reads. 2) We extracted genomic DNA sequences 20 nt downstream of the end of each cluster. If the number of adenosine nucleotides was more than 65% in the genomic sequence, we ignored the corresponding cluster and marked it as caused by mispriming during the second strand synthesis in the RT reaction. 3) We ignored clusters overlapping with other clusters in the same orientation by 2 nt or less. 4) We attached clusters to the closest gene in the same orientation. If no gene could be identified within 2,000 nt, the cluster was ignored. 5) In cases with multiple 3’-UTR isoforms identified, we calculated the frequency of occurrence for each isoform and ignored isoforms occurring at a frequency of less than 1% independently from the number of reads that form this cluster. The logo plots used to visualize our results were produced using the WebLogo 3 suite (Crooks et al. 2004).

### Extraction of 3’-UTR regions from the *C. elegans* genome

The 3’-UTRs used in the experiments described in **Figure 5C and Supplemental Figure S11-S13** were initially cloned from N2 *wild type C. elegans* genomic DNA using PCR with Platinum Taq Polymerase (Invitrogen). Genomic DNA template was prepared as previously described (Blazie et al. 2017). Forward DNA primers were designed to include approximately 30 nucleotides upstream of the translation STOP codon and include the endogenous translation STOP codon. We used the Gateway BP Clonase II Enzyme Mix (Invitrogen) to clone the 3’-UTR region into Gateway entry vectors. The DNA primer was modified to include the attB Gateway recombination elements required for insertion into pDONR P2RP3 (Invitrogen). The reverse DNA primers were designed to end between 200 and 250 nucleotides downstream of the RNA cleavage site and to include the reverse recombination element attB for cloning into pDONR P2RP3 (Invitrogen). At the conclusion of the recombination step, the entry vectors containing the cloned 3’-UTR regions were transformed into Top10 competent cells (Thermo Fisher Scientific), using agar plates containing 20mg/μL of kanamycin. The plasmids were then recovered, and clones were confirmed using Sanger sequencing with the M13F primer. The list of primers used in this study is available in **Supplemental Table S2**.

### Data Access

Strains and plasmids are available upon request. The authors affirm that all data necessary for confirming the conclusions of the article are present within the article, figures and supplemental figures, and tables and supplemental tables. The results of our analyses are available in WormBase (www.WormBase.org) (Lee et al. 2018) and in our 3’-UTR-centric website www.UTRome.org.

## Acknowledgements

We thank Heather Hrach for insights and review of the manuscript. We thank Gabrielle Richardson for maintaining the *C. elegans* strains used in this manuscript. This work was supported by the National Institute of Health grant number 1R01GM118796.

## Author Contribution

HSS and MM designed the experiments. MM developed and executed the bioinformatic analysis and 3’-UTR cluster preparation. HSS performed the rescue experiments in **Figure 5 and Supplemental Figure S11-S13**. PLC performed the homology modeling in **Figure 4B and Supplemental Figure S10** and helped write the manuscript. CG assisted with the experiments and performed the analysis in **Supplemental Figure S1**. SO contributed to the experiments in **Supplemental Figures S11-S13**. MM uploaded the results to the WormBase and UTRome.org database. MM and HSS led the analysis and interpretation of the data, assembled the Figures, and wrote the manuscript. All authors read and approved the final manuscript.

