## Supplemental Figures for "The *C. elegans* 3’-UTRome V2: an updated genomic resource to study 3’-UTR biology"

Additional File 1 for:

#### This PDF includes:

- **Figures S1-S14**
- **Supplemental Materials and Methods**

**Table of Contents**

### Additional File 1


**Figure S1:** Protein alignment of members of the CPC complex in five species ……………………………………………………..….3

**Figure S2:** Results of the RNAi experiments of the *C. elegans* CPC...…………14

**Figure S3**: Bioinformatic pipeline used in this study …….……………………..15

**Figure S4:** Comparison with Tourasse et al., 2017……………………………….16

**Figure S5:** PAS site usage in genes with multiple 3'-UTR isoforms.……………..17

**Figure S6:** Detection of the PAS element in genes lacking a canonical AAUAAA hexamer.…………………………......……18

**Figure S7:** GO term analysis for genes with 1, 2 or 3 3'-UTR isoforms ………...19

**Figure S8:** Detection of the 'UGUA' element in *C. elegans* 3'-UTRs……………20

**Figure S9:** Detection of enriched elements in 3'-UTRs of genes with 2 3'-UTR isoforms………………………………………21

**Figure S10:** Nucleotide binding site of the human CPSF160-WDR33-CPSF30 complex ……………………..…………22

**Figure S11:** *In vivo* cleavage assay for M03A1.3 ……………………………….23

**Figure S12:** *In vivo* cleavage assay for Y106G6H.9…………………...…………24

**Figure S13**: *In vivo* cleavage assay for *ges-1* ………………..……...………..25

**Figure S14:** miRNA target analysis in genes with 2 3'-UTR isoforms which either gain or lose a miRNA binding site……...……...……26

**Supplemental Materials and Methods:** ……………………...……...……27

#### CPSF-1 (CPSF160)

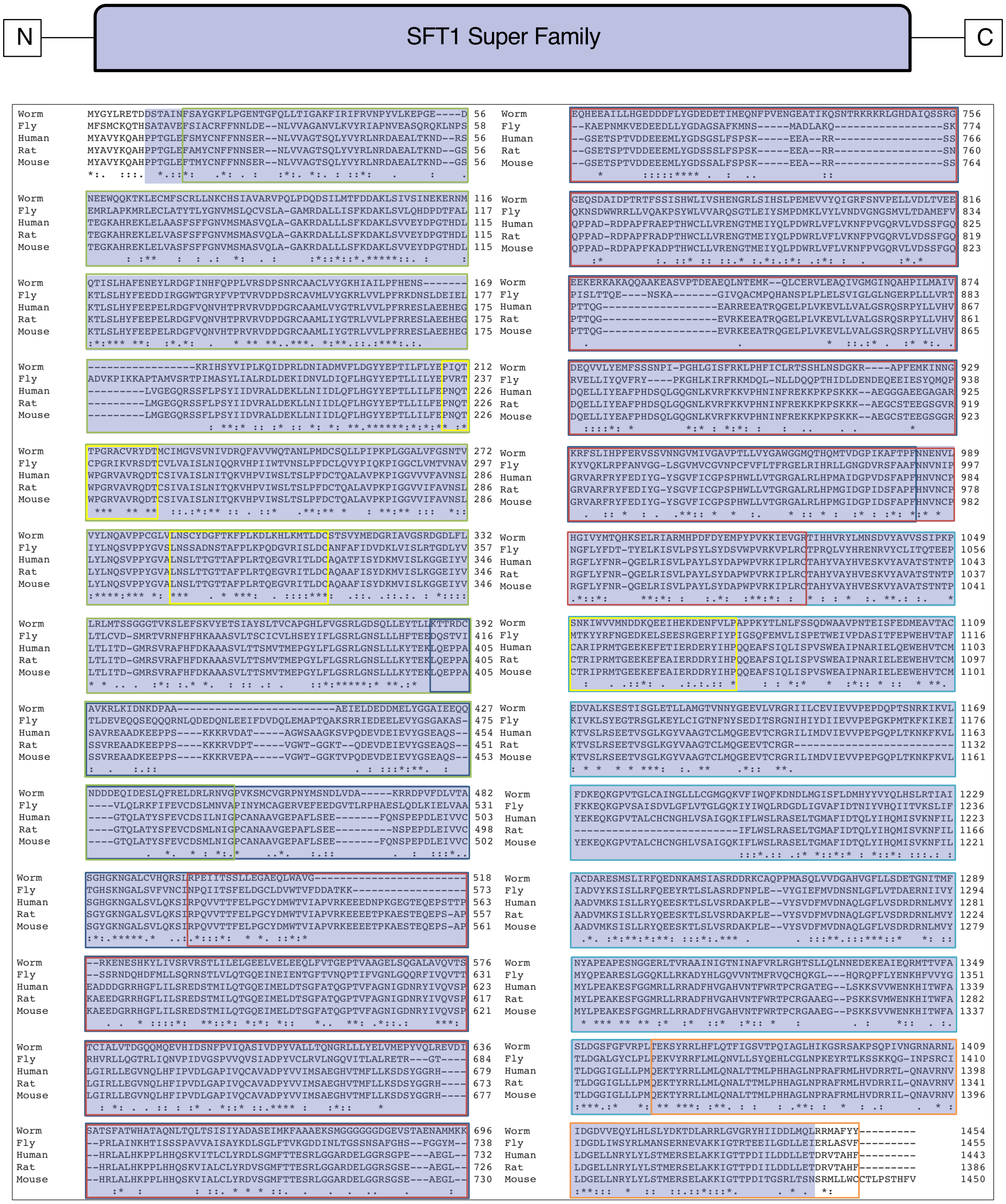


- Residues 400 to 973 interact with HAT-C domain of CstF-77 (Bai et al., 2007)
- bP1 domain made of WD40 repeats (Clerici et al., 2017, Sun et al., 2018)
- bP2 domain made of WD40 repeats (Clerici et al., 2017, Sun et al., 2018)
- bP3 domain made of WD40 repeats interacts with CPSF30 (Clerici et al., 2017, Sun et al., 2018)
- C-terminal domain (Clerici et al., 2017, Sun et al., 2018)
- Elongated loops 1-3 stabilize interaction with Wdr33 NTD (Clerici et al., 2017, Sun et al., 2018)

**Supplemental Figure S1: Protein alignment of members of the CPC complex in five organisms.** Amino acid sequence alignment for the members of the CPC from five organisms produced with Clustal Omega Multiple Sequence Alignment. Conserved domains produced with the Batch Conserved Domain Search on NCBI are represented by the highlighted regions of each figure. Known domains determined from previously published literature are outlined.

### CPSF Complex

#### CPSF-2 (CPSF100)

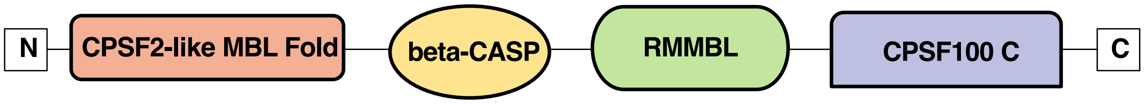


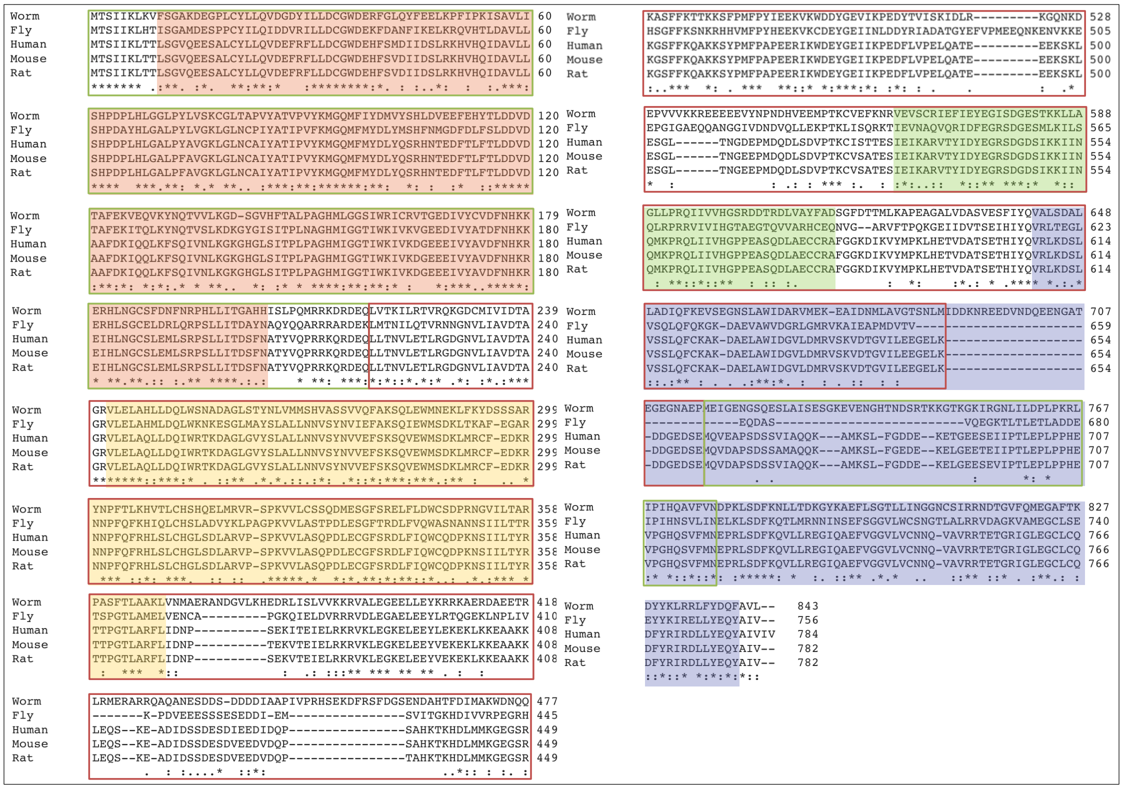


Metallo-beta-lactamase domain (Mandel et al., 2006)

Beta-CASP domain (Mandel et al., 2006)

#### CPSF-3 (CPSF73)

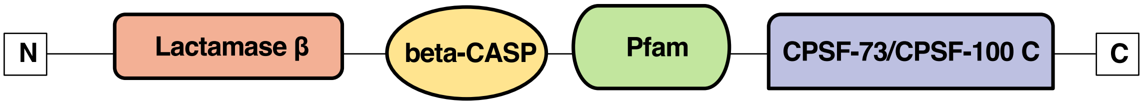


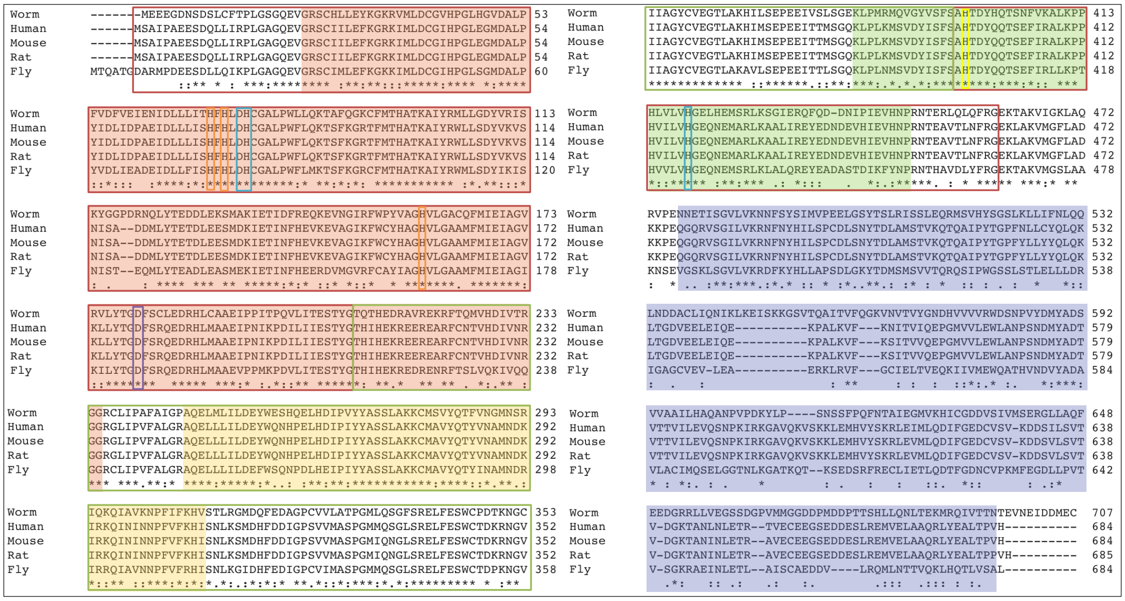


Metallo-beta-lactamase domain (Mandel et al., 2006)

Beta-CASP domain (Mandel et al., 2006)

Bridging ligand to Zinc atoms (Mandel et al., 2006)

Ligands to Zn1 (Mandel et al., 2006)

Ligands to Zn2 (Mandel et al., 2006)

General acid for catalysis (Mandel et al., 2006)

#### CPSF-4 (CPSF30)

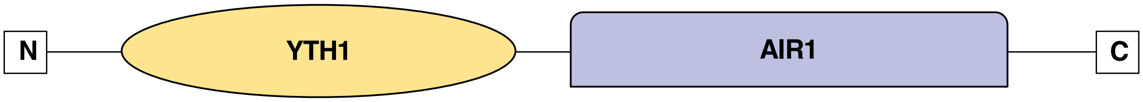


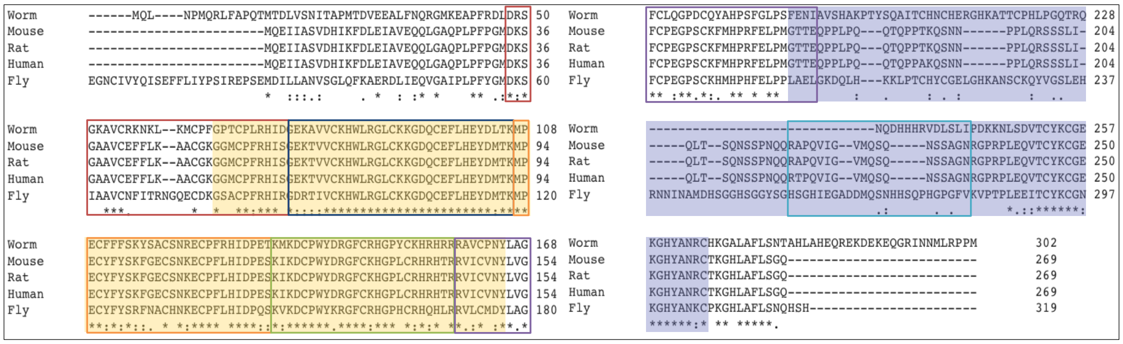

Zinc Finger 1 (Clerici et al., 2017)
Zinc Finger 2, contacts A1 and A2 of PAS (Clerici et al., 2017, Sjun et al., 2018)
Zinc Finger 3, contacts A4 and A5 of PAS (Clerici et al., 2017, Sjun et al., 2018)
} Contact Wdr33 and CPSF160
Zinc Finger 4 (Clerici et al., 2017)
Zinc Finger 5 (Clerici et al., 2017)
} Contact Fip1
Zinc Knuckle (Clerici et al., 2017)

#### FIPP-1 (FIP1)

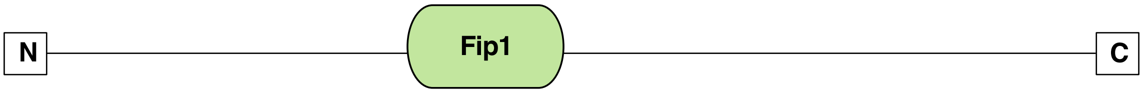


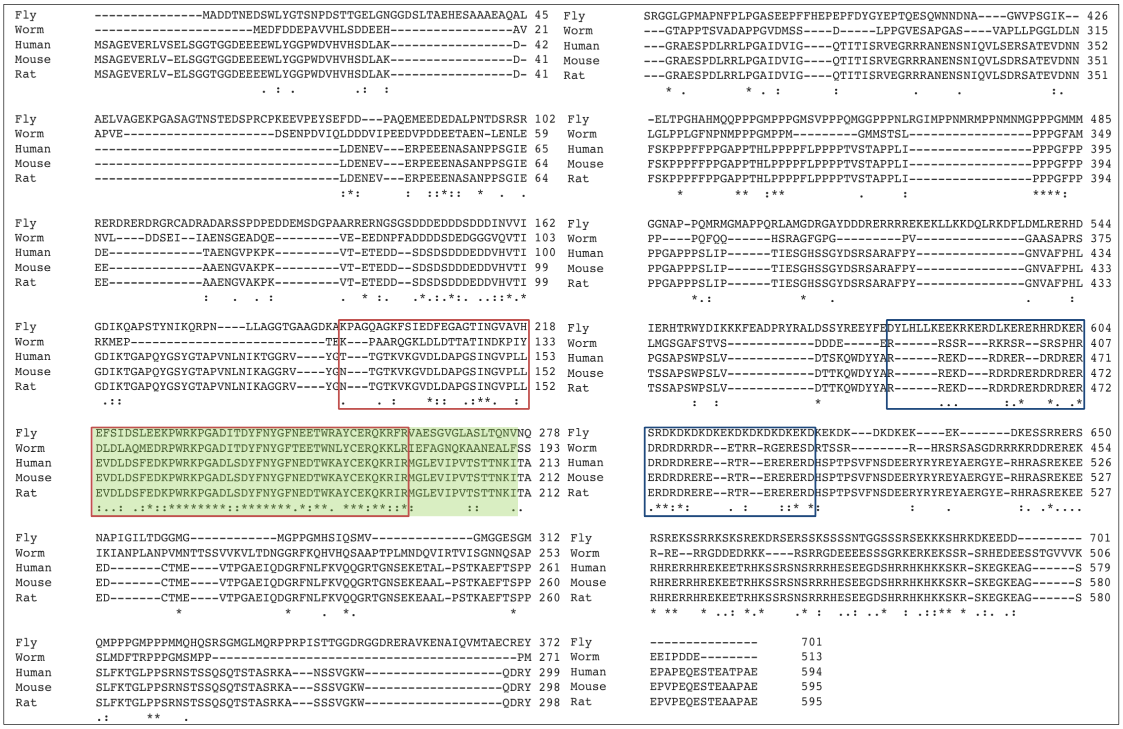

Conserved Domain contacts CPSF30 (Clerici et al., 2017)
RE/D region interacts with CFIm68/59 (Zhu et al., 2018)

#### PFS-2 (WDR33)

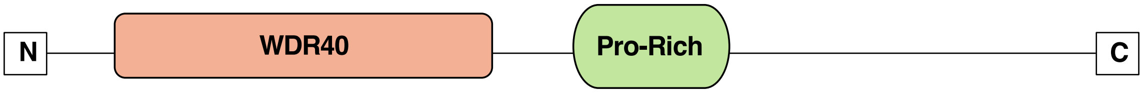


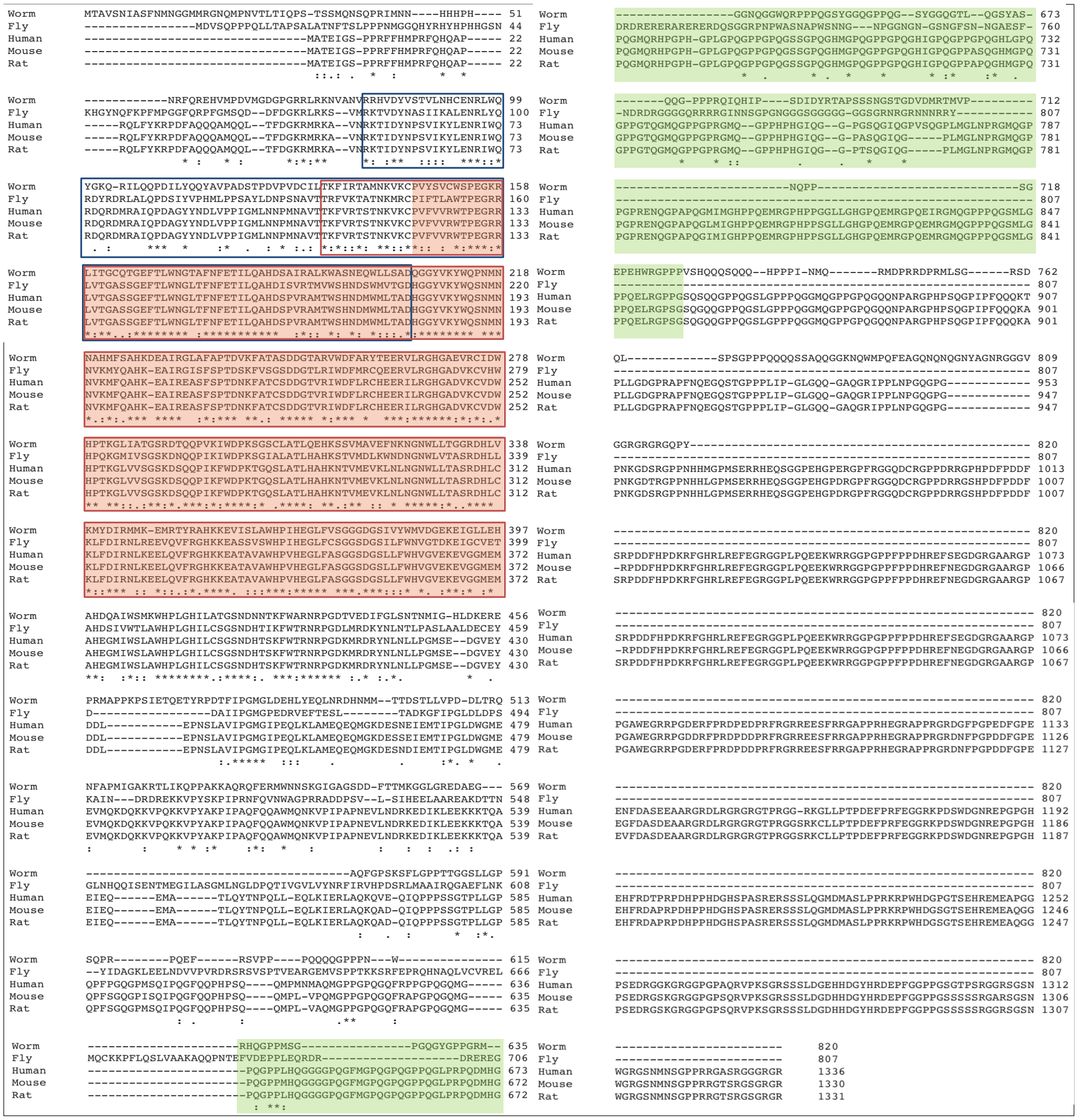


N-terminal domain, contacts CPSF160 and CPSF30 (Clerici et al., 2017)

WD40 repeats, contact U3 and A6 of PAS (Clerici et al., 2017)

CstF Complex

#### CPF-1 (CstF-50)

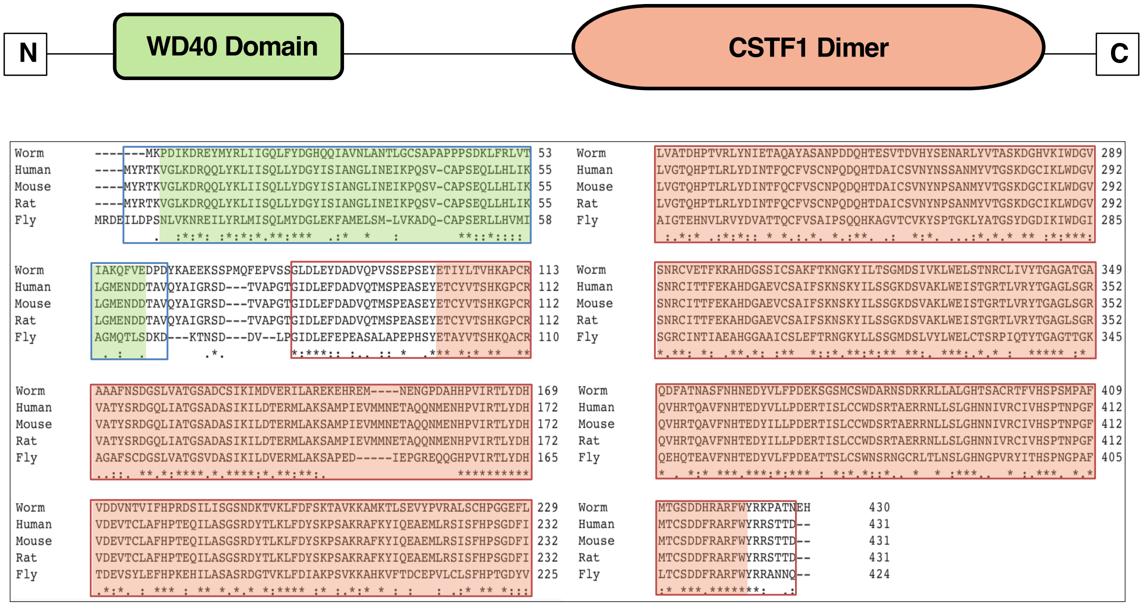


WD40 Repeats (Yang et al., 2018)

N-terminal domain (Yang et al., 2018)

#### CPF-2 (CstF-64)

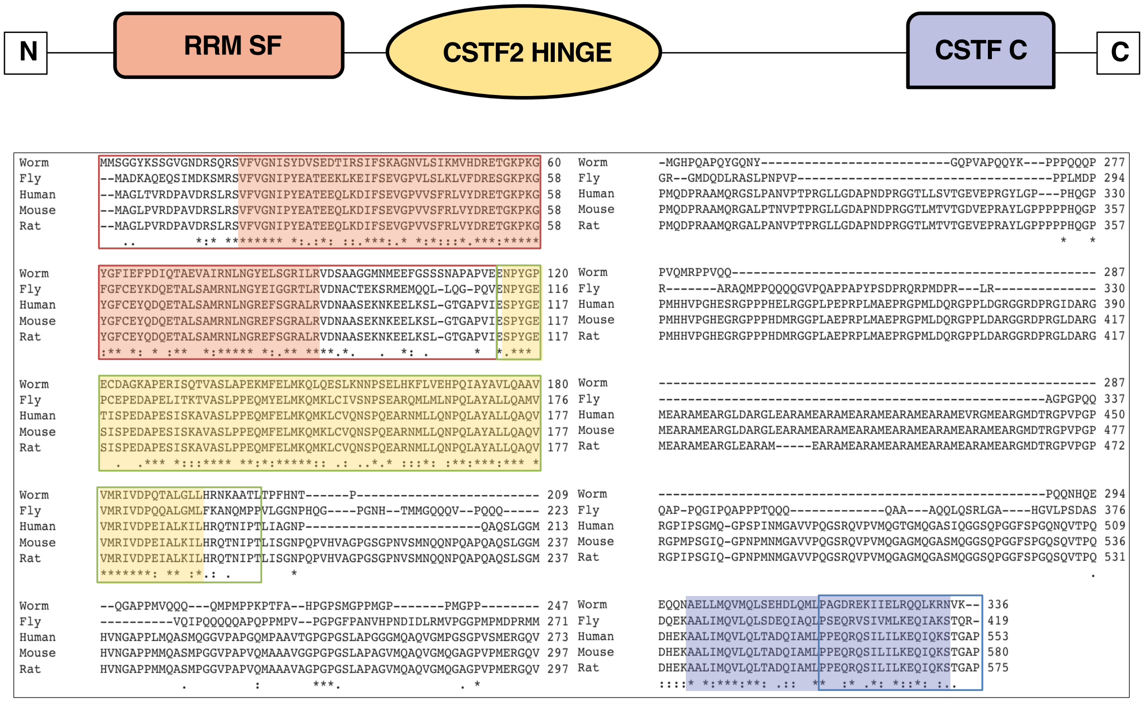


RRM domain (Yang et al., 2018)

HINGE domain (Yang et al., 2018)

C-terminal domain (Yang et al., 2018)

### CstF Complex

#### SUF-1 (CstF77)

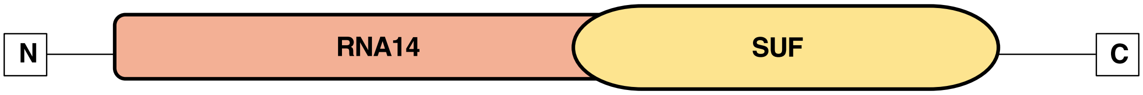


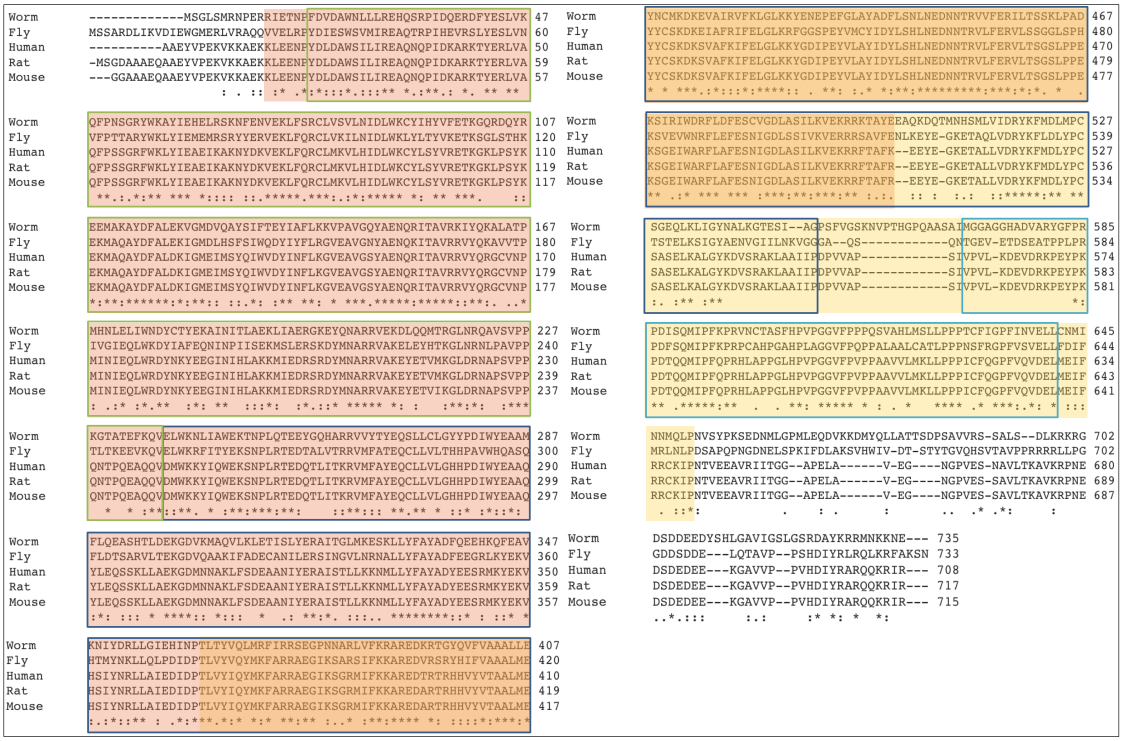

Proline-rich segment that facilitates interactions with CstF-64 and CstF-55 (Bai et al., 2007)

HAT-N Domain (Bai et al., 2007)

HAT-C Domain that interacts with CPSF160 (Bai et al., 2007)

CFIm Complex

#### CFIM-1 (CFIm25)

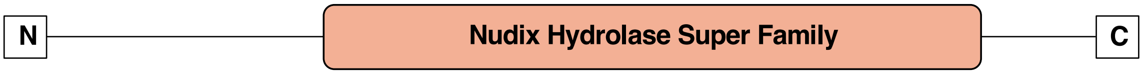


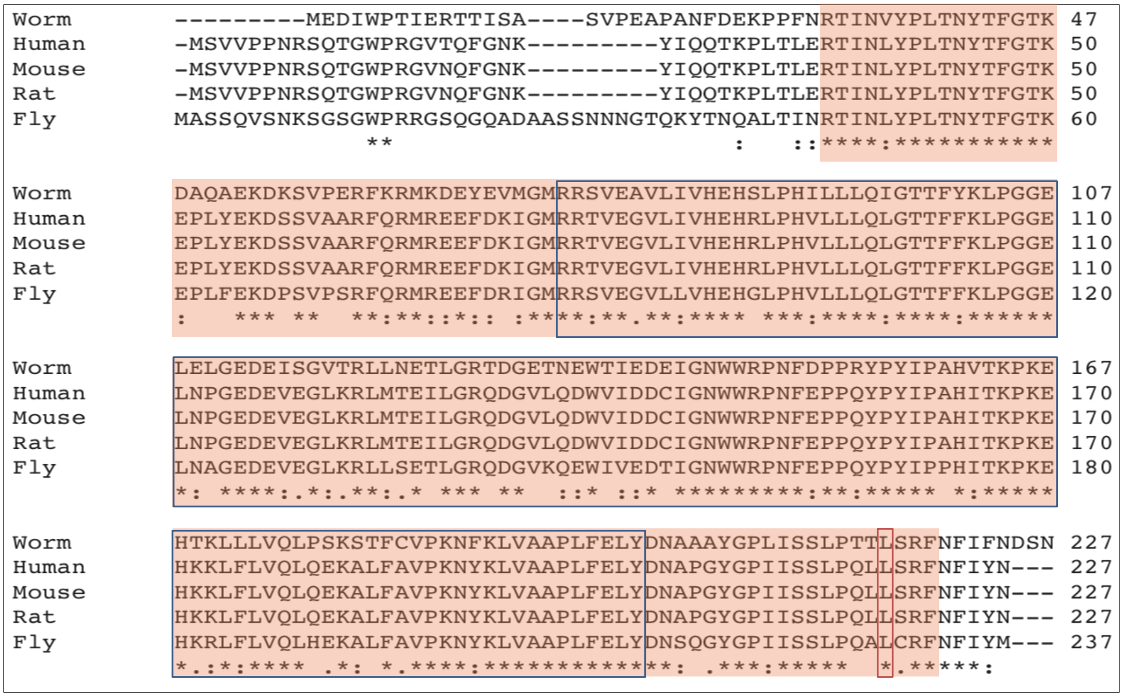


Nudix domain (Yang et al., 2011) CFIm68 and CFIm59 tethering site (Zhu et al.,2017)

#### CFIM-2 (CFIm68)

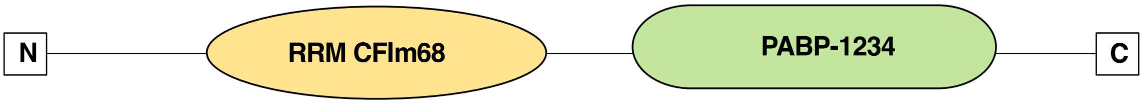


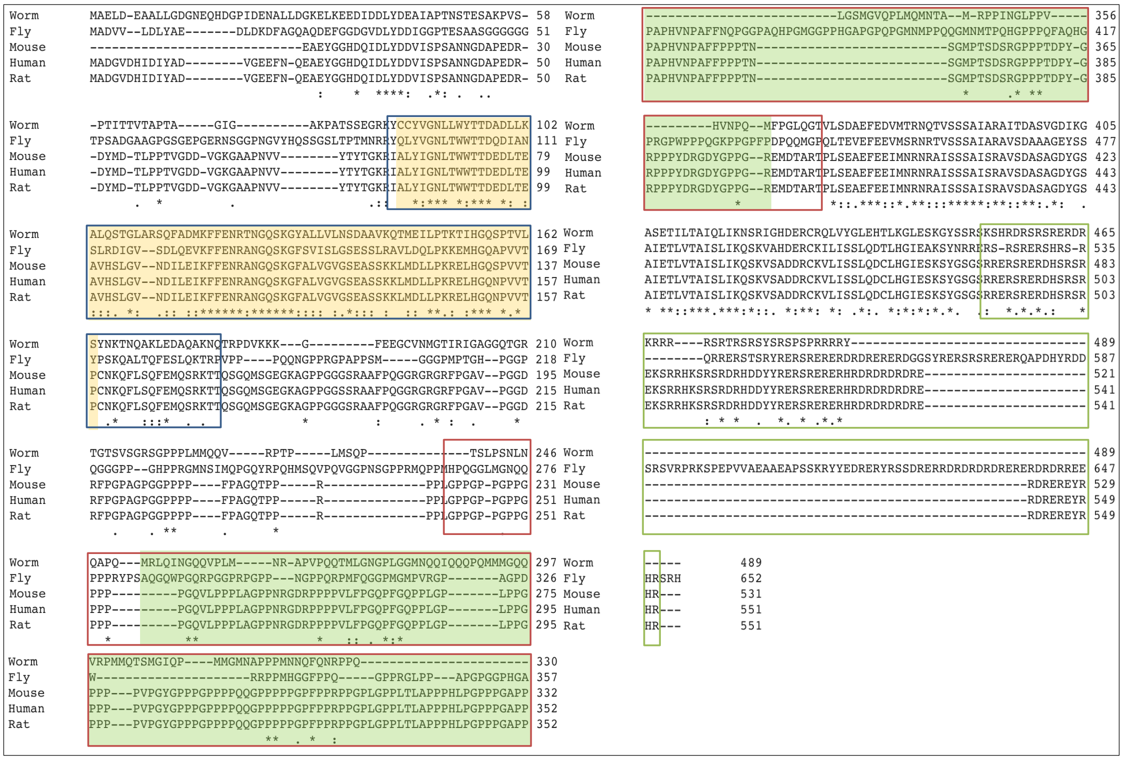


RRM domain (Yang et al., 2011, Martin et al., 2010) RS/RD/RE region (Yang et al., 2011)

Pro/Gly-rich region (Yang et al., 2011)

**CLPF-1 (CLP1)**

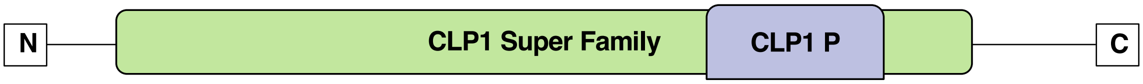


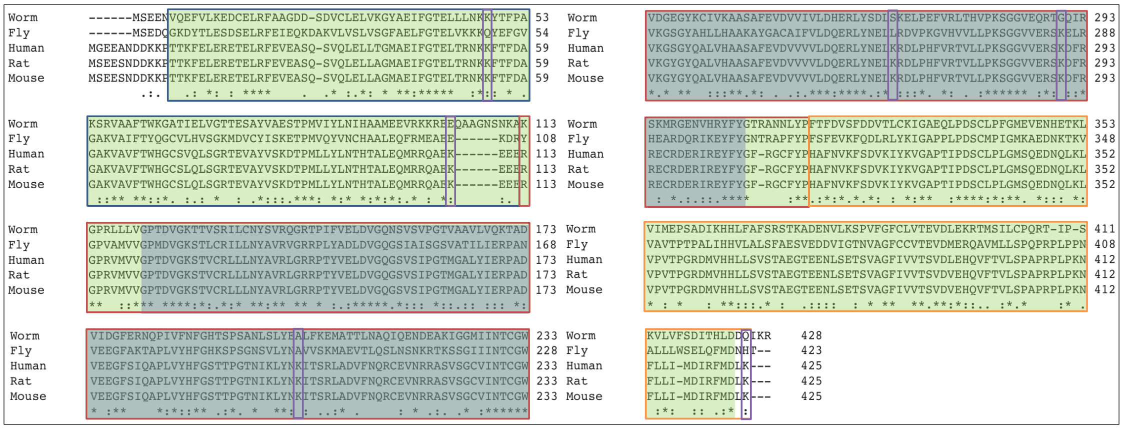

N-terminal domain (Dikfidan et al., 2014)

Polynucleotide kinase domain (Dikfidan et al., 2014)

C-terminal domain (Dikfidan et al., 2014)

Residues that crosslink to PCF-11 (Schäfer et al., 2018)

#### RBPL-1 (RBBP6)

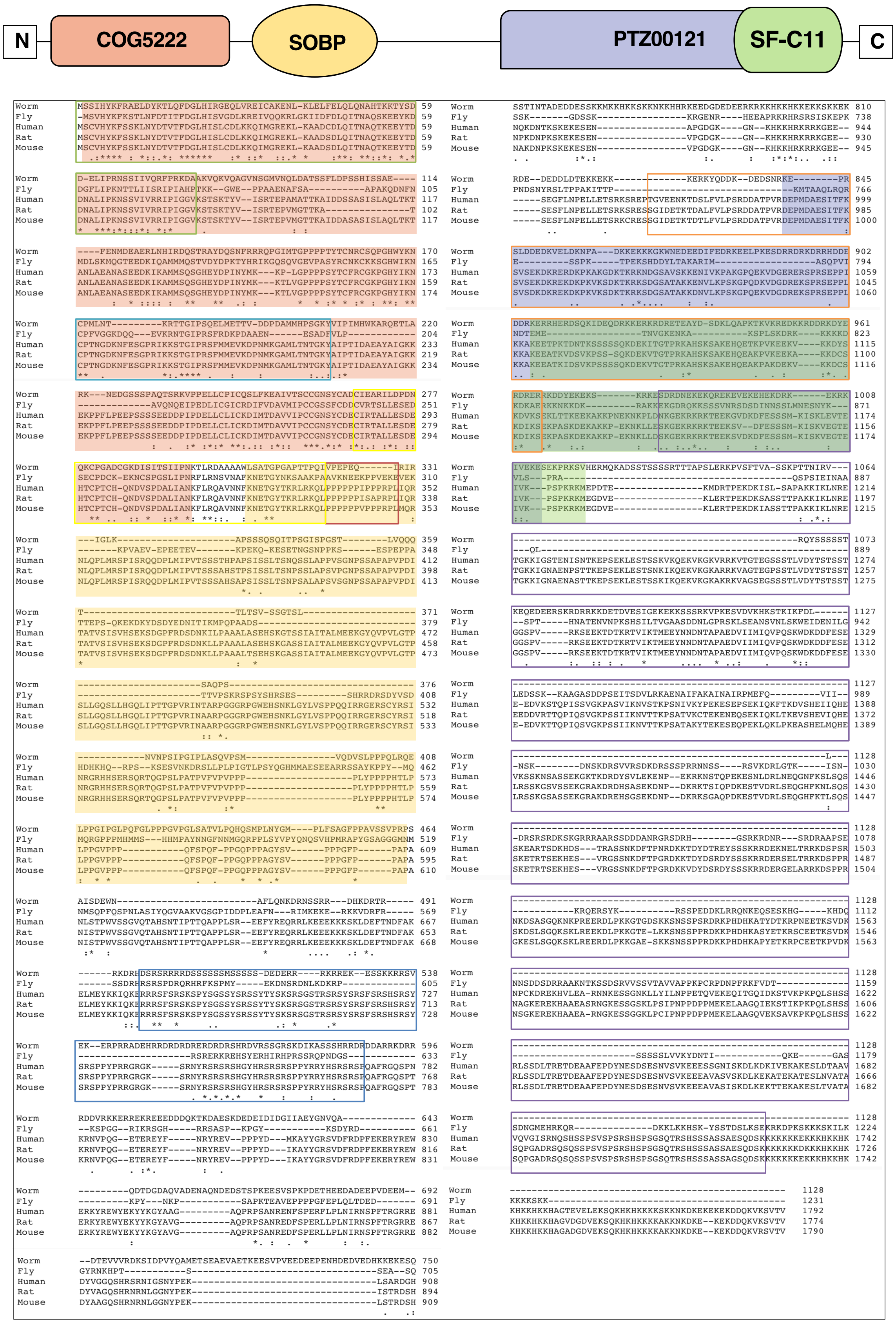


- Domain With No Name (DWNN), ubiquitin-like (Pugh et al., 2006)
- RING finger domain (Chibi et al., 2008, Lee et al., 2014)
- Proline-rich domain (Pugh et al., 2006)
- SR domain (Pugh et al., 2006)
- Rb-binding domain (Pugh et al., 2006)
- p53-binding domain (Pugh et al., 2006)
- Zinc knuckle (Lee et al., 2014)

#### PAP-1 (PAP1N)

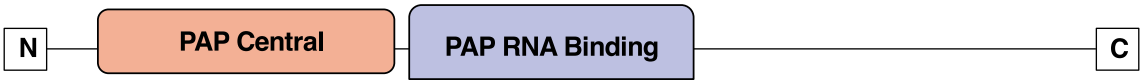


```
Worm     ----------------MSATEKDKTPLLGVSQPISLAHPDSKDIAQTTLLIETLKKFGSY 44
Fly      MWNSEPTHRQHHQHNGNSTSGGPPAKQLGMTSAISLAEPRPEDLQRTDELRGSLEPYNVF 60
Human    --------MPFPVTTQGSQQTQPPQKHYGITSPISLAAPKETDCVLTQKLIETLKPFGVF 52
Mouse    --------MPFPVTTQGSQQTQPPQRHYGITSPISLAAPKETDCLLTQKLIETLKPFGVF 52
Rat      --------MPFPVTTQGSQQTQPPQRHYGITSPISLAAPKETDCLLTQKLIETLKPFGVF 52
                                 *           *::. **** *    *    *   *   :*: :. :

Worm     EPKEETEQRMEVLRNLNRLVKEWVKNVTAM-KIPNGEGVNAGGKLFTFGSYRLGVHSSGA 103
Fly      ESQDELNHRMEILAKLNTLVKQWVKEISVSKNMPESAAEKLGGKIYTFGSYRLGVHHKGA 120
Human    EEEEELQRRILILGKLNNLVKEWIREISESKNLPQSVIENVGGKIFTFGSYRLGVHTKGA 112
Mouse    EEEEELQRRILILGKLNNLVKEWIREISESKNLPQSVIENVGGKIFTFGSYRLGVHTKGA 112
Rat      EEEEELQRRILILGKLNNLVKEWIREISESKNLPQSVIENVGGKIFTFGSYRLGVHTKGA 112
         * ::* ::*: :* :** ***:*:::::   ::*:.   : ***::********** .**

Worm     DIDTLAVVPRHIDRSDFFTSFKEMLNNDPNVTELHGVEEAFVPVMKLKYSGVELDILFAR 163
Fly      DIDALCVAPRNIERTDYFQSFFEVLKKQPEVTECRSVEEAFVPVIKMNFDGIEIDLLFAR 180
Human    DIDALCVAPRHVDRSDFFTSFYDKLKLQEEVKDLRAVEEAFVPVIKLCFDGIEIDILFAR 172
Mouse    DIDALCVAPRHVDRSDFFTSFYDKLKLQEEVKDLRAVEEAFVPVIKLCFDGIEIDILFAR 172
Rat      DIDALCVAPRHVDRSDFFTSFYDKLKLQEEVKDLRAVEEAFVPVIKLCFDGIEIDILFAR 172
         ***:*.*.**:::*:*:* ** : *: : :*.: :.********:*: :.*:*:*:****

Worm     LALKEVPDTQELSDDNLLRNLDQESVRSLNGCRVAEQLLKLVPRQKEFCVTLRAIKLWAK 223
Fly      LSLKEIPDDFDLRDDNLLRNLDHRSVRSLNGCRVTDEILALVPNIENFRLALRTIKLWAK 240
Human    LALQTIPEDLDLRDDSLLKNLDIRCIRSLNGCRVTDEILHLVPNIDNFRLTLRAIKLWAK 232
Mouse    LALQTIPEDLDLRDDSLLKNLDIRCIRSLNGCRVTDEILHLVPNIDNFRLTLRAIKLWAK 232
Rat      LALQTIPEDLDLRDDSLLKNLDIRCIRSLNGCRVTDEILHLVPNIDNFRLTLRAIKLWAK 232
         *:*: :*:  :* **.**:*** ..:********::::* ***. .:* ::**:******

Worm     NHGIYSNSMGFFGGITWAILVARACQLYPNASPSRLVHRMFFIFSTWTWPHPVVLNEMNN 283
Fly      KHGIYSNSLGYFGGVTWAMLVARTCQLYPNAAAATLVHKFFLVFSRWKWPNPVLLKHPDN 300
Human    RHNIYSNILGFLGGVSWAMLVARTCQLYPNAIASTLVHKFFLVFSKWEWPNPVLLKQPEE 292
Mouse    RHNIYSNILGFLGGVSWAMLVARTCQLYPNAIASTLVHKFFLVFSKWEWPNPVLLKQPEE 292
Rat      RHNIYSNILGFLGGVSWAMLVARTCQLYPNAIASTLVHKFFLVFSKWEWPNPVLLKQPEE 292
         .*.**** :*::**::**:****:*******  : ***::*::** * **:**:*:. ::

Worm     DRNDIPTLCELVWDPRRKNTDRFHVMPIITPAFPEQNSTHNVTRSTATVIKNEICEALEI 343
Fly      V-----NLRFQVWDPRVNASDRYHLMPIITPAYPQQNSTFNVSESTKKVILTEFNRGMNI 355
Human    C-----NLNLPVWDPRVNPSDRYHLMPIITPAYPQQNSTYNVSVSTRMVMVEEFKQGLAI 347
Mouse    C-----NLNLPVWDPRVNPSDRYHLMPIITPAYPQQNSTYNVSVSTRMVMVEEFKQGLAI 347
Rat      C-----NLNLPVWDPRVNPSDRYHLMPIITPAYPQQNSTYNVSVSTRMVMVEEFKQGLAI 347
               .*   ***** : :**:*:*******:*:****.**: **  *:  *: ..: *

Worm     CRDISEGKSKWTALFEEVNFFSRYKHFIALIMAAPNEEEELNYGGFLESRIRLLVQSLER 403
Fly      TDEIMLGRIPWERLFEAPSFFYRYRHFIVLLVNSQTADDHLEWCGLVESKVRLLIGNLER 415
Human    TDEILLSKAEWSKLFEAPNFFQKYKHYIVLLASAPTEKQRLEWVGLVESKIRILVGSLEK 407
Mouse    TDEILLSKAEWSKLFEAPNFFQKYKHYIVLLASAPTEKQRLEWVGLVESKIRILVGSLEK 407
Rat      TDEILLSKAEWSKLFEAPNFFQKYKHYIVLLASAPTEKQRLEWVGLVESKIRILVGSLEK 407
           :*  .:  *  ***  .** :*:*:*.*:  : . .:.*:: *::**::*:*: .**:

Worm     NQDIIIAHNDPNKHKPSPNAKF--------------------DVNPENKRVTVWFIGLEF 443
Fly      NPHIALAHVNPKCFEFKKGQSANNSQNNSGNEDDLKQSQGNQSAVTSAPFCSMWFIGLEF 475
Human    NEFITLAHVNPQSFPAPKENP------------------------DKEEFRTMWVIGLVF 443
Mouse    NEFITLAHVNPQSFPAPKESP------------------------DREEFRTMWVIGLVF 443
Rat      NEFITLAHVNPQSFPAPKESP------------------------DREEFRTMWVIGLVF 443
         *  * :** :*: .                                ::*.*** *

Worm     AEHA----KTLDLTNEIQRFKTNVELQASNVKGIGPNCQVQIDMFYVKRNSLIQVISAAD 499
Fly      ER---SENLNVDLTESIQNFTEHVMMHGVNIKMLKEG--MTIDARHVKRKQLSLYLDSDF 530
Human    KKTENSENLSVDLTYDIQSFTDTVYRQAINSKMFEVD--MKIAAMHVKRKQLHQLLPNHV 501
Mouse    KKTENSENLSVDLTYDIQSFTDTVYRQAINSKMFELD--MKIAAMHVKRKQLHQLLPSHV 501
Rat      KKTENSENLSVDLTYDIQSFTDTVYRQAINSKMFELD--MKIAAMHVKRKQLHQLLPSHV 501
          .       .:*** .** *.   *   :. * * :  .  : *   :***:.*    :

Worm     LRRGRRWKKVVPIATNTSVSSSTPRSVVRTTSTSSVPTTPTGLAAPKTPLSASVSATNEP 559
Fly      LKRERKSMESHNNFNNTLLAN-RKRLSTELAQSQ-DPLPP-GQQ-------PSSGN---- 576
Human    LQKKKKHSTEGVKLT--ALNDSSLDLSMDSDNSMSVPSPT-SAT-KTSPLNSSGSS---- 553
Mouse    LQKRKKHSTEGVKLT--ALNDSSLDLSMDSDNSMSVPSPT-SAM-KTSPLNSSGSS---- 553
Rat      LQKRKKHSTEGVKLA--ALNDSSLDLSMDSDNSMSVPSPT-SAM-KTSPLNSSGSS---- 553
         *:: ::             : .          .:   *    .          * .

Worm     DSTTNGTPLSRKR-SMDEES---------STTVTSQISDESVPKKKTRDDTLEENRVSMV 609
Fly      RGRDSGAKIQRLSDSLTEENSNASSDMGAGTPTTPTTAQLSAPSFKSSGKNGSE------ 630
Human    QGRNSPAPAV-----------------TAASVTNIQATEVSVPQVNSSESSG-------- 588
Mouse    QGRNSPAPAV-----------------TAASVTSIQASEVSVPQANSSESPG-------- 588
Rat      QGRNSPAPAV-----------------TAASVTSIQASEVSVPQANSSESPG-------- 588
          .  . :                      .: ..   :: *.*. ::  .

Worm     VEVSNVVVEQRTKVVQEIVDLQADNGLNTSNGLEASEQKMEVPQSV-------------- 655
Fly      -----------IDVVEQEPTQPHNNGNASSN-----TTTTEVACSXQPPTLHPHPPHQQP 674
Human    --------------------------GTSS---------ESIPQTATQP-------AISP 606
Mouse    --------------------------GPSS---------ESIPQTATQP-------AISP 606
Rat      --------------------------GPSS---------ESIPQTATQP-------AISP 606
                                   :*           .:   :

Worm     ------------------------------------------------------------ 655
Fly      PPAPPQPPAHHHH---HQKFRKNNNAAAAAASNSIFNQRSILHAASSLSAALSITGHKRK 731
Human    PPKPTVSRVVSSTRLVNPPPRSSGNAATSG--NA----------AT--KIPTPIVGVKRT 652
Mouse    PPKPTVSRVVSSTRLVNPSPRPSGN--------T----------AT--KVPNPIVGVKRT 646
Rat      PPKPTVSRVVSSTRLVNPSPRPSGN--------A----------AT--KVPNPIVGVKRT 646

Worm     ------------------------------------------------------------ 655
Fly      QQTATSTTMISSSNIAHSNYTQHSTHNNNGGGGGSHYNKTYYIDDDDEDNNQPLLTPNNQ 791
Human    SSPHK-------------EESPKKTKT-----------EEEQLDTET---------STTQ 679
Mouse    SSPNK-------------EESPKKTKT-----------EEEQVDLET---------SAVQ 673
Rat      SSPHK-------------EESPKKTKT-----------EEEQVGMET---------SAAP 673

Worm     -----------------------------------------------       655
Fly      TQQQQKAVRITA---TSASPATSVPVAASAAPPAPTISL---------       827
Human    SETIQTAASLLASQKTSSTDLSDIP--ALPANPIPVIKNSIKLRLNR       724
Mouse    SETVPASASLLASQKTSSTDLSDIP--ALPANPIPVIKNSIKLRLNR       718
Rat      SEAAPASASLVASQKTSSTDLSDIP--ALPANPIPVIKNSIKLRLNR       718
```

#### SYMK-1 (Symplekin)

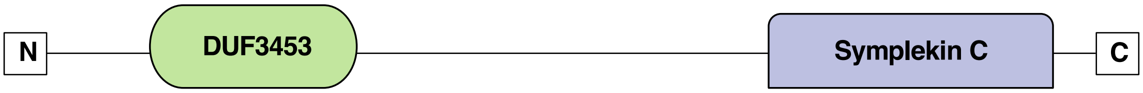


```
Worm      MDYIQG------------------LNEENETASERIGEALKEARDAETIEKKLLSLSTAM 42
Fly       MDSIIGRSQFVS-ETANLFTDEK-----TATARAKVVDWCNELVIASP-STKCELLAKVQ 53
Human     MASGSGDSVTRRSVASQFFTQEEGPGIDGMTTSERVVDLLNQAALITN-DSKITVLKQVQ 59
Mouse     MASSSGDSVTRRSVASQFFTQEEGPSIDGMTTSERVVDLLNQAALITN-DSKITVLKQVQ 59
Rat       MASGSGDSVTRRSVASQFFTQEEGPGIDGMTTSERVVDLLNQAALITN-DSKITVLKQVQ 59
          *    *                     *:  :: :  ::       ..*   *  .

Worm      HLLIDPSLSISILDNFLTEMLEFAELNDSRILCLLVDFLLKASAKDFTLCNKTVERYSFY 102
Fly       ETVL--GSCAELAEEFLESVLSLAHDSNMEVRKQVVAFVEQVCKVKVELLPHVINVVSML 111
Human     ELII--NKDPTLLDNFLDEIIAFQADKSIEVRKFVIGFIEEACKRDIELLLKLIANLNML 117
Mouse     ELII--NKDPTLLDNFLDEIIAFQADKSIEVRKFVIGFIEEACKRDIELLLKLIANLNML 117
Rat       ELII--NKDPTLLDNFLDEIIAFQADKSTEVRKFVIGFIEEACKRDIELLLKLIANLNML 117
          . ::  .    : ::** .:: :   .. .:   :: *: :..  .. *  : :   .:

Worm      LIPNKSIKRYESVIKRVVVASTNLYPIVLEFA--IMDKNDNAESCWDAFNLLKNRICMLV 160
Fly       LRD-----NSAQVIKRVIQACGSIYKNGLQYLCSLMEPGDSAEQAWNILSLIKAQILDMI 166
Human     LRD-----ENVNVVKKAILTMTQLYKVALQWMVKSRVISELQEACWDMVSAMAGDIILLL 172
Mouse     LRD-----ENVNVVKKAILTMTQLYKVALQWMVKSRVISDLQEACWDMVSSMAGEIILLL 172
Rat       LRD-----ENVNVVKKAILTMTQLYKVALQWMVKSRVISDLQEACWDMVSSMAAEIILLL 172
          *       .  .*:*:.: :  .:*   *::       .:  * .*: .. :   *  ::

Worm      SDDHEGVRTVTVKFLEALILCQSPKPRELATGSNISWAREANTRFNRISLSDVPRSHRFL 220
Fly       DNENDGIRTNAIKFLEGVVVLQSFADED-----------SLKRDGDFSLADVPDHCTLF 214
Human     DSDNDGIRTHAIKFVEGLIVTLSPRMADSE----I-----PRRQEHDISLDRIPRDHPYI 223
Mouse     DSDNDGIRTHAIKFVEGLIVTLSPRMADSE----V-----PRRQEHDISLDRIPRDHPYI 223
Rat       DSDNDGIRTHAIKFVEGLIVTLSPRMADSE----V-----PRRQEHDISLDRIPRDHPYI 223
          ..:::*:** ::**:*.:::  *    :               :   :**  :*      :

Worm      SYHKTQLEAEENFSALLKQTTVAEATSQNLITVIESLCMITRCRPQWENALPRVFDVIKA 280
Fly       RREKLQEEGNNILDILLQFHGTTHISSVNLIACTSSLCTIAKMRPIFMGAV---VEAFKQ 271
Human     QYNVLWEEGKAALEQLLKFMVHPAISSINLTTALGSLANIARQRPMFMSEV---IQAYET 280
Mouse     QYNVLWEEGKAAVEQLLKFMVHPAISSINLTTALGSLANIARQRPMFMSEV---IQAYET 280
Rat       QYNVLWEEGKAALEQLLKFMVHPAISSINLTTALGSLANIARQRPMFMSEV---IQAYET 280
            .    *.:  .. **:        :* ** :   **. *:: ** : . :   .:. :

Worm      LHSNVPPMLSKGQVKFLRKSFKYNLLRFLKLPASVPLQQKITTMLTNYLGASPREVQQSI 340
Fly       LNANLPPTLTDSQVSSVRKSLKMQLQTLLKNRGAFEFASTIRGMLVD-LGSSTNEIQKLI 330
Human     LHANLPPTLAKSQVSSVRKNLKLHLLSVLKHPASLEFQAQITTLLVD-LGTPQAEIARNM 339
Mouse     LHANLPPTLAKSQVSSVRKNLKLHLLSVLKHPASLEFQAQITTLLVD-LGTPQAEIARNM 339
Rat       LHANLPPTLAKSQVSSVRKNLKLHLLSVLKHPASLEFQAQITTLLVD-LGTPQAEIARNM 339
          *::*:** *:..**. :**.:* :*  .**  .:. :   *  :*.: **:   *: : :

Worm      PPELIQKIAPPRP--PQHPAEPVAKRPKIQNQIFEDDDDDDDEAGPSTSTV-NAKDARTE 397
Fly       PKMDKQEMARRQKRILENAAQSLAKRARLACE-QQDQQQREMELDTEE-----LERQKQK 384
Human     PSSKDTRKRPRDD-----SDSTL-KKMKLEPNLGEDDEDKDLEPGPSGTSKASAQISGQS 393
Mouse     PSSKDSRKRPRDD-----TDSTL-KKMKLEPNLGEDDEDKDLEPGPSGTSKASAQISGQS 393
Rat       PSSKDSRKRPRDD-----TESTL-KKMKLEPNLGEDDEDKDLEPGPSGTSKASAQISGQS 393
          *     .               . : *: ::  :  :*::: : * . .        :    .

Worm      AIDMTAKFI-MECLNHETVMNLVKISLYTLPSEMPAAFASSYTPIANAGTEPNRQELSEL 456
Fly       STRVNEKFLAEHFRNPETVVTLVLEFLPSLPTEVPQKFLQEYTPIREMSIQQQVTNISRF 444
Human     DTDITAEFL-QPLLTPDNVANLVLISMVYLPEAMPASFQAIYTPVESAGTEAQIKHLARL 452
Mouse     DTDITAEFL-QPLLTPDNVANLVLISMVYLPETMPASFQAIYTPVESAGTEAQIKHLARL 452
Rat       DTDITAEFL-QPLLTPDNVANLVLISMVYLPETMPASFQAIYTPVESAGTEAQIKHLARL 452
             :. :*:     . :.* .**   :  **  :*  *   ***: . . : :  .::.:

Worm      MAVQMTNKEIGPGYEWLQQQRKKEYEARNKARSEGMAIAQ-----------------TP 498
Fly       FGEQLSEKRLGPGAATFSRE--PPMRVKKVQAIESTLTAME--VDEDA---------VQK 491
Human     MATQMTAAGLGPGVEQTKQCKEEPKEEKVV-KTESVLIKRRLSAQGQAISVVGSLSSMSP 511
Mouse     MATQMTAAGLGPGVEQTKQCKEEPKEEKVV-KPESVLIKRRLSVQGQAISVVGSQSTMSP 511
Rat       MATQMTAAGLGPGVEQTKQCKEEPKEEKVV-KPESVLIKRRLSVQGQAISVVGSQSTMSP 511
          :. *::   :***    .:      . :     *.

Worm      IHEPN-MSNRVPAQIVKQSLQE-INTLPVI---QKAKKAFNLVEEAVVFDDKEAAEMFEL 553
Fly       LSEEEFQRKEEATKKLRETMERAKGEQTVIEKMKERAKTLKLQEITKPLPRNLKEKFLTD 551
Human     LEEEAPQAKRRPEPII-------PVTQPRLAGAGGRKKIFRLSDVLKPLTDAQVEAMKLG 564
Mouse     LEEEVPQAKRRPEPII-------PVTQPRLAGAGGRKKIFRLSDVLKPLTDAQVEAMKLG 564
Rat       LEEEVPQAKRRPEPII-------PVTQPRLAGAGGRKKIFRLSDVLKPLTDAQVEAMKLG 564
          : *     :.     :              :       * :.* :    :       :

Worm      AYESVLQAERRVVAGGARLMYQKLVVRLTTRFWEDCTPFEEKLIEFVLADHKKRNDLALL 613
Fly       AVRRILNSERQCIKGGVSSKRRKLVTVIAATFPD---NVRYGIMEFILEDIKQRIDLAFS 608
Human     AVKRILRAEKAVACSGAAQVRIKILASLVTQFNS---GLKAEVLSFILEDVRARLDLAFA 621
Mouse     AVKRILRAEKAVACSGAAQVRIKILASLVTQFDS---GFKAEVLSFILEDVRARLDLAFA 621
Rat       AVKRILRAEKAVACSGAAQVRIKILASLVTQFDS---GFKAEVLSFILEDVRARLDLAFA 621
          * . :*.:*:    .*.     *::. :.: * .    ..  ::.*:* * : * ***:

Worm      WLCELYAQYQGYSNCALFMKEMIAGQEGLTQAQRLDRYDQAMCKMLDAMLERNMEKEAL- 672
Fly       WLFEEYSLLQGFTRHTYVK----------TENRPDHAYNELLNKLIFGIGERCDHKDKII 658
Human     WLYQEYNAYLAAGA-----------------SGSLDKYEDCLIRLLSGLQEKPDQKDGI- 663
Mouse     WLYQEYNAYLAAGT-----------------SGTLDKYEDCLICLLSGLQEKPDQKDGI- 663
Rat       WLYQEYNAYLAAGP-----------------SGTLDKYEDCLICLLSGLQEKPDQKDGI- 663
          ** : *    .                     . *:: :  :: .: *:  .*: :

Worm      -FYKVLLETPLLTPNAIERLKQVCLAKE-NEHGMAMLRELIMTRNRQRPQLLQFLFGLFF 730
Fly       LIRRVYLEAPILPEVSIGHLVQLSLDDEFSQHGLELIKDLAVLRPPRKNRFVRVLLNFSV 718
Human     -FTKVVLEAPLITESALEVVRKYCEDESRTYLGMSTLRDLIFKRPSRQFQYLHVLLDLSS 722
Mouse     -FTKVVLEAPLITESALEVIRKYCEDESRAYLGMSTLGDLIFKRPSRQFQYLHVLLDLSS 722
Rat       -FTKVVLEAPLITESALEVIRKYCEDESRAYLGMSTLGDLIFKRPSRQFQYLHVLLDLSS 722
           : :* **:*::   ::  :  : .  ..    *:  : :* . *  :: : ::.*:.:

Worm      MERPELRSSCLEVVKELCYLPF-IRSSLSDQARMQIHDCLQESPPMYMRSS-----EDSD 784
Fly       HERLDLRDLAQAHLVSLYHVHKILPARIDEFALEWLKFIEQESPPAAVFSQDFGRPTEEP 778
Human     HEKDKVRSQALLFIKRMYEKE-QLREYVEKFALNYLQLLVHPNPPSVLFGADKDTE-VAA 780
Mouse     HEKDRVRSQALLFIKRMYEKE-QLREYVEKFALNYLQLLVHPNPPSVLFGADKDTE-VAA 780
Rat       HEKDRVRSQALLFIKRMYEKE-QLREYVEKFALNYLQLLVHPNPPSVLFGADKDTE-VAA 780
           *:  :*. .   :  :      :   :.. *  * ::   : .**  :  .

Worm      QWTDEMYKNSLAVYSTLMPSDPL-LLIPLASVYAQSTNVFKRVVLRSLEPVFRQLSQ--E 841
Fly       DWREDTTKVCFGLAFTLLPYKPEVYLQQICQVFVSTSAELKRTILRSLDIPIKKMGVESP 838
Human     PWTEETVKQCLYLYLALLPQNHK-LIHELAAVYTEAIADIKRTVLRVIEQPIRGMGMNSP 839
Mouse     PWTEETVKQCLYLYLALLPQNHK-LIHELAAVYTEAIADIKRTVLRVIEQPIRGMGMNSP 839
Rat       PWTEETVKQCLYLYLALLPQNHK-LIHELAAVYTEAIADIKRTVLRVIEQPIRGMGMNSP 839
           * ::  * .: :  :*:* .     :  :. *:..  :  :**.:**:  ::  :: :.

Worm      MVISLIEDCPYGAETLVARLVVLLTERIT-PSTDLIQKLKILHDERKMDIRALLPIIGGL 900
Fly       TLLQLIEDCPKGMETLVIRIIYILTERVPSPHEELVRRVRDLYQNKVKDVRVMIPVLSGL 898
Human     ELLLLVENCPKGAETLVTRCLHSLTDKVP-PSPELVKRVRDLYHKRLPDVRFLIPVLNGL 898
Mouse     ELLLLVENCPKGAETLVTRCLHSLTDKVP-PSPELVKRVRDLYHKRLPDVRFLIPVLNGL 898
Rat       ELLLLVENCPKGAETLVTRCLHSLTDKVP-PSPELVKRVRDLYHKRLPDVRFLIPVLNGL 898
           :: *:*:** * **** * :  **:::  *    :*::::: *:.::  *:* ::*::.**

Worm      EREEVVRLIPTFIFRAEYQKSVNVLFRKLYTVRDPQ--TGNLVFDPIEVIKEYHKIEPKN 958
Fly       TRSELISVLPKLIKLNPA--VVKEVFNRLLGIGAEFAHQ-TMAMTPTDILVALHTIDTSV 955
Human     EKKEVIQALPKLIKLNPI--VVKEVFNRLLGTQHGEGNSALSPLNPGELLIALHNIDSVK 956
Mouse     EKKEVIQALPKLIKLNPI--VVKEVFNRLLGTQHGEGNSALSPLNPGELLIALHNIDSVK 956
Rat       EKKEVIQALPKLIKLNPI--VVKEVFNRLLGTQHGEGNSALSPLNPGELLIALHNIDSVK 956
           :.*::  :*.:*         *: :*.:*             : * ::   *.*:

Worm      DNEAELLVNNLEFLFDPALLKPDTASQAIEAVFKWENVPFLFLHSLYTLFHKFKTFESFV 1018
Fly       CDIKAIVKATSLCLAERDLYTQEVLMAVLQQLVEVTPLPTLMMRTTIQSLTLYPRLANFV 1015
Human     CDMKSIIKATNLCFAERNVYTSEVLAVVMQQLMEQSPLPMLLMRTVIQSLTMYPRLGGFV 1016
Mouse     CDMKSIIKATNLCFAERNVYTSEVLAVVMQQLMEQSPLPMLLMRTVIQSLTMYPRLGGFV 1016
Rat       CDMKSIIKATNLCFAERNVYTSEVLAVVMQQLMEQSPLPMLLMRTVIQSLTMYPRLGGFV 1016
           :   ::  .   : :  : .  .:: .::  :*  :* *:::    :   :  : .**

Worm      ANLFYKVTEKKMYQQSDRWKQAFFKCIKELKTKAYPAVITFLS------FEEYEELKEVL 1072
Fly       MNLLQRLIIKQVWRQKVIWE-GFLKTVQRLKPQSMPILLHLPPAQLVDALQQCPDLRPAL 1074
Human     MNILSRLIMKQVWKYPKVWE-GFIKCCQRTKPQSFQVILQLPPQQLGAVFDKCPELREPL 1075
Mouse     MNILARLIMKQVWKYPKVWE-GFIKCCQRTKPQSFQVILQLPPQQLGAVFDKCPELREPL 1075
Rat       MNILARLIMKQVWKYPKVWE-GFIKCCQRTKPQSFQVILQLPPQQLGAVFDKCPELREPL 1075
           *:: ::  *::::       *: .*:*  :. * ::   :: :          :::  :*:  *

Worm      G-------------DGIVAEFKIIYSTLATQQQK-----N--MDEKIKEELHDKERENR- 1111
Fly       SEYAESMQDEPMNGSGITQQVLDIISGKSVDVFVTDESGGYISAEHIKKEAPDPSEISVI 1134
Human     LAHVRSFTPH--QQAHIPNSIMTILEASGKQ-----------EP-EAKEAPAGPLEEDD- 1120
Mouse     LAHVRSFTPH--QQAHIPNSIMTILEATGKQ-----------EP-EVKEAPSGPLEEDD- 1120
Rat       LAHVRSFTPH--QQAHIPNSIMTILEASGKQ-----------EP-EVKEAPSGSLEEDD- 1120
                      *  ..  * .  . :                 . *:   .  . .

Worm      -----------------------------ERDKRLRREEKKEKEREKE-RTRESGKERS 1140
Fly       STVPVLTSLVPLPVPPPIGSDLNQPLPPGED----------------------------- 1165
Human     -LEPLTLA----------P--APAPRPPQDLIGLRLAQEKALKRQLEEEQKLKPGGVGAP 1167
Mouse     -LEPLALALAPAPAPAPAP--APAPRPPQDLIGLRLAQEKALKRQLEEEQKQKPTGIGAP 1177
Rat       -LEPLALALAPALA----P--APAPRPPQDLIGLRLAQEKALKRQLEEEQKQKPTGVGAP 1173

Worm      SRR--------------------------------------------------------- 1143
Fly       ------------------------------------------------------------ 1165
Human     SS---SSPSPSPSARPGPPPSEEAMDFREEGPECETPGIFISMDDDSGLTEAALLDSSLE 1224
Mouse     AACVSSTPSVPAAARAGPTPAEEVMEYREEGPECETPAIFISMDDDSGLAETTLLDSSLE 1237
Rat       TSSVSSTPLVGPAARAGPTPAEEVMEYREEGPECETPAIFISMDDDSGLAETTLLDSSLE 1233

Worm      -----------------------------------------------------  1143
Fly       -----------------------------------------------------  1165
Human     GPLPKETAAGGLTLKEERSPQTLAPVGEDAMKTPSPAAEDAREPEAKGNS     1274
Mouse     GPLPKEAAAVGSSSKDERSPQNLSHAVEEALKTSSPE---TREPESKGNS     1284
Rat       GPLPKEAAAVGPSSKDERSPQNLSHAVEEALKTSSPE---AREPESKGNS     1280
```

| Name | Experiment # | Eggs/Hatched | Lethality (%) |
|---|---|---|---|
| ***cpsf-1*** *(CPSF160)* | 1 | 163/11 | **93.7** |
| | 2 | 238/5 | **97.9** |
| | 3 | 149/1 | **99.3** |
| ***cpsf-2*** *(CPSF100)* | 1 | 68/37 | **64.8** |
| | 2 | 178/28 | **86.4** |
| | 3 | 221/25 | **89.8** |
| ***cpsf-4*** *(CPSF30)* | 1 | 323/5 | **98.5** |
| | 2 | 699/17 | **97.6** |
| | 3 | 204/3 | **98.6** |
| ***cpf-1*** *(CstF-50)* | 1 | 251/76 | **76.8** |
| | 2 | 200/64 | **75.8** |
| | 3 | 185/25 | **88.1** |
| ***cpf-2*** *(CstF-64)* | 1 | 446/54 | **89.2** |
| | 2 | 138/14 | **90.8** |
| | 3 | 61/3 | **95.3** |
| ***cfim-1*** *(CFIm25)* | 1 | 249/23 | **91.5** |
| | 2 | 196/20 | **90.7** |
| | 3 | 154/2 | **98.7** |
| ***cfim-2*** *(CFIm68)* | 1 | 137/1 | **99.3** |
| | 2 | 282/27 | **91.3** |
| | 3 | 260/32 | **89.0** |
| ***lrp-2*** *(CFIm59)* | 1 | 311/116 | **72.8** |
| | 2 | 281/97 | **74.3** |
| | 3 | 222/93 | **70.5** |
| ***symk-1*** *(Symplekin)* | 1 | 62/4 | **93.9** |
| | 2 | 19/4 | **82.6** |
| | 3 | 114/5 | **95.8** |
| ***rbpl-1*** *(RBBP6)* | 1 | 293/0 | **100** |
| | 2 | 344/0 | **100** |
| | 3 | 116/0 | **100** |
| ***pcf-11*** *(PCF11)* | 1 | 141/5 | **96.6** |
| | 2 | 105/0 | **100** |
| | 3 | 172/5 | **97.2** |
| ***clpf-1*** *(CLP1)* | 1 | 286/23 | **92.6** |
| | 2 | 289/18 | **94.1** |
| | 3 | 208/17 | **92.4** |
| ***pkc-3*** *(negative control)* | 1 | 5/0 | **100** |
| | 2 | 8/0 | **100** |
| | 3 | 18/1 | **94.7** |
| | 4 | 51/1 | **98.1** |
| | 5 | 3/0 | **100** |
| | 6 | 22/0 | **100** |
| | 7 | 53/0 | **100** |
| | 8 | 6/0 | **100** |
| | 9 | 13/0 | **100** |
| | 10 | 19/0 | **100** |
| | 11 | 5/0 | **100** |
| | 12 | 4/0 | **100** |

**Supplemental Figure S2: Results of the RNAi experiments of the *C. elegans* CPC.** Twelve genes for the members of the *C. elegans* CPC were knocked-down using RNAi. Clones/rows are color-coded as from Figure 1. The human orthologs of each gene are shown in parenthesis in the first column. For each RNAi experiment we use 15 worms, and the number of eggs unhatched vs hatched at the end of the experiment were counted. The percent lethality was consistently high across all tested clones. *pkc-3* RNAi was used as a negative RNAi control, since it is known to induce strong embryonic lethality.

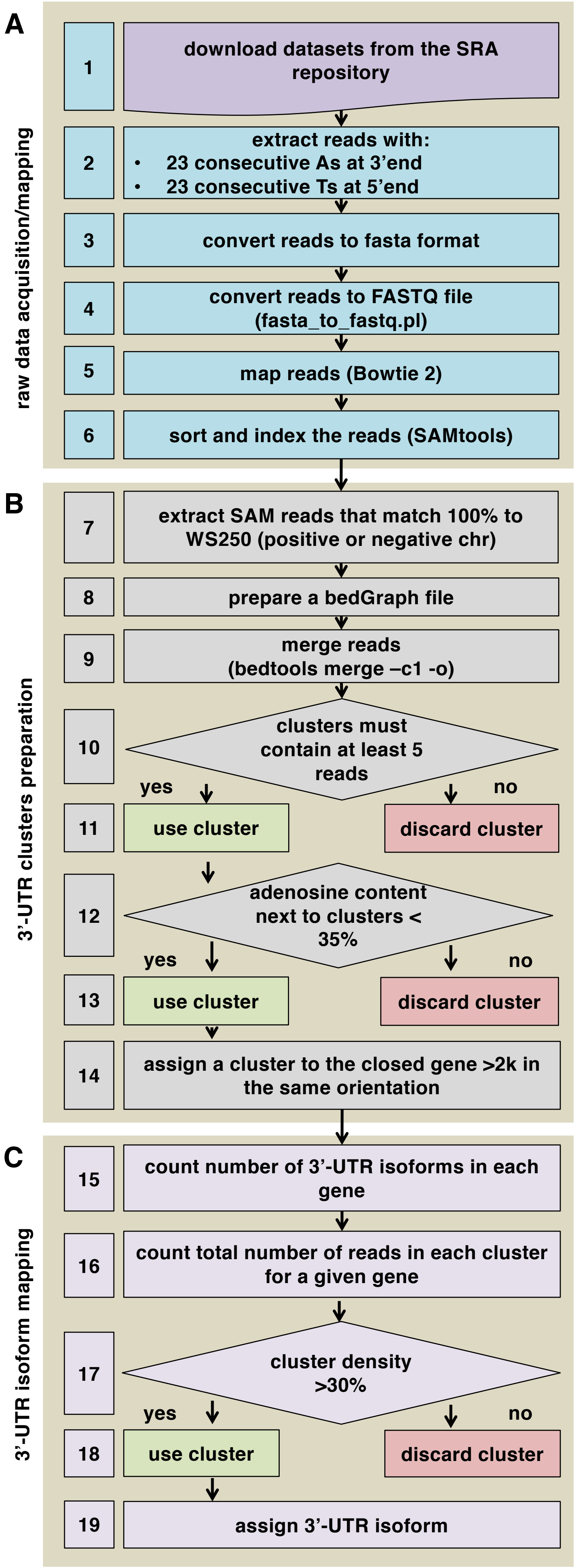


**Supplemental Figure S3: Bioinformatic Pipeline used in this study.** The pipeline uses raw transcriptome datasets downloaded from the public repository SRA trace archive to extract and map 3'-UTR end clusters to the closest protein-coding genes in the correct orientation.
The pipeline is divided in three large steps: A) Acquisition/Mapping, B) 3'-UTR cluster preparation and C) 3'-UTR isoforms mapping.
In the acquisition/mapping step, we used custom made Perl scripts to extract reads with 23 consecutive As at the 3'-end or 23 consecutive Ts at the 5'-end and then mapped these filtered reads to the WS250 version of the *C. elegans* genome (Bowtie 2). We then sorted and indexed the reads for visualization purposes.
In the 3'-UTR cluster preparation step, we extracted SAM reads with 100% match to the WS250 and used them to prepare a new bedGraph file (BEDTools).
We then merged the reads and discarded the clusters with less than 5 reads.
Restrictive parameters for cluster identification and 3'-UTR end mapping included the discard of clusters with an adenosine content of <35% downstream of its end.
Clusters were assigned to a mapped 3'-UTR end and attached to the closest gene with 2,000 nt in the same orientation.
At the completion of these steps we performed the 3'-UTR isoform mapping step, which consists of the counting and assignment the total number of 3'-UTR isoforms to a given gene.
We discarded clusters with a density of less than 30% of the total number of reads.

# A

| mismatch (+/-) | stringent | raw |
|---|---|---|
| | | n=27,264 |
| 1 | 1,604 | 2,167 |
| 5 | 2,637 | 3,373 |
| 10 | 2,898 | 3,842 |
| 20 | 3,702 | 5,437 |

| mismatch (+/-) | Mangone et al., 2010 | Jan et al., 2011 |
|---|---|---|
| | | n=27,264 |
| 1 | 930 | 285 |
| 5 | 1,717 | 989 |
| 10 | 2,229 | 1,664 |
| 20 | 3,436 | 3,106 |

# B

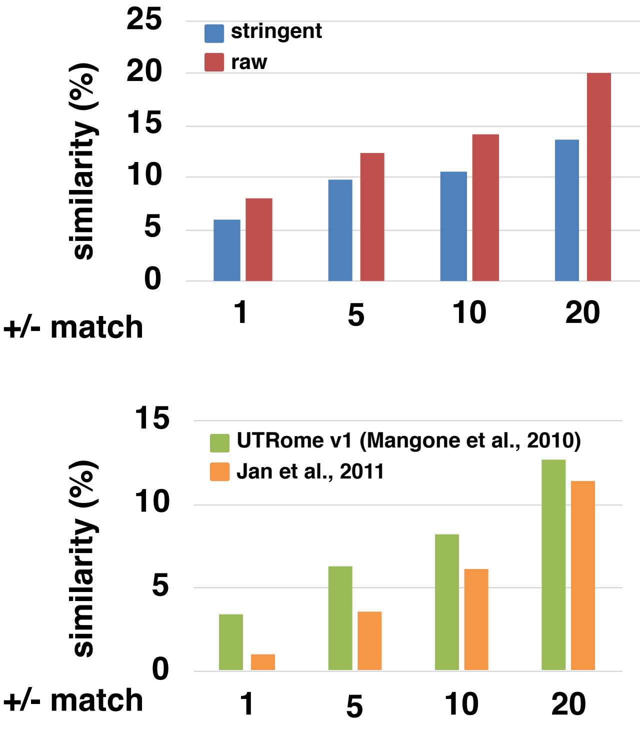


# C

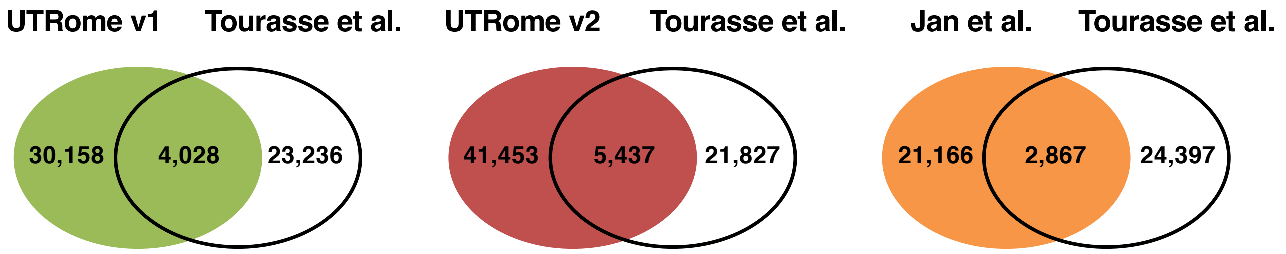


**Supplemental Figure S4: Comparison with Tourasse et al., 2017.** We have downloaded the poly(A) sites mapped in Tourasse et al. and performed a comparison with the 3'-UTRs present in our 3'-UTRome v2. A) Top Panel: number of mapped poly(A) sites from Tourasse et al. that match our stringent and raw datasets within +/-1 nt, +/- 5 nt, +/- 10 nt or +/- 20 nt. Bottom Panel: number of poly(A) sites in common between Tourasse et al. and Mangone et al. and between Tourasse et al. and Jan et al. within +/-1nt, +/- 5nt, +/- 10nt or +/- 20nt. B) Bar chart showing the % of similarity of the two datasets in Panel A. C. Venn diagrams comparing the 3'-UTRs shared (+/- 20nt) between Tourasse et al., and the UTRome v1 (Mangone et al., - (green), this study (UTRome v2 - red), and Jan et al. (orange). We have used our unfiltered dataset to compare UTRome v2 with Tourasse et al.

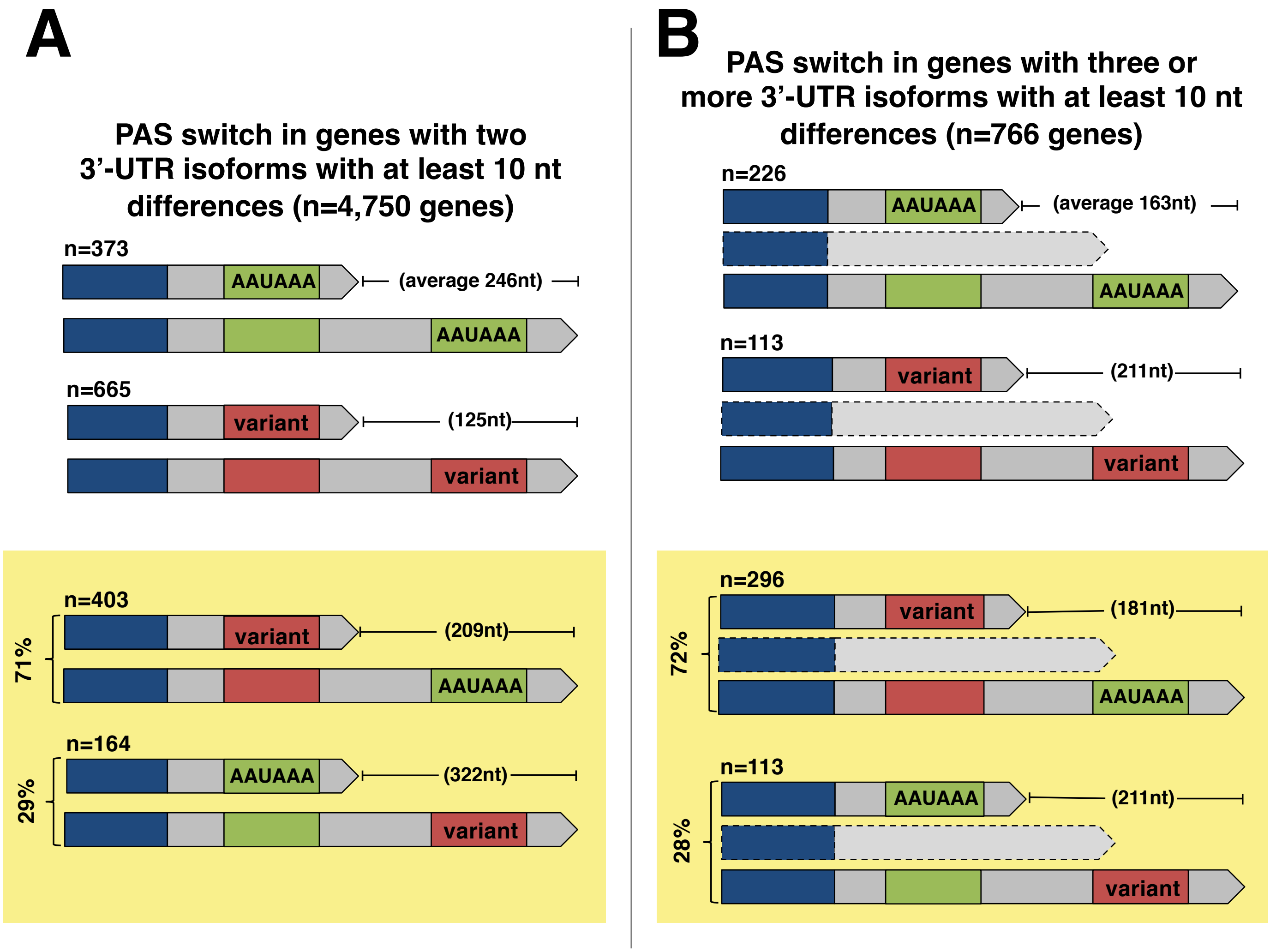


**Supplemental Figure S5: PAS site usage in genes with multiple 3'-UTR isoforms.** A) In genes with only two 3'-UTR isoforms with a difference of at least 10 nt between isoforms, 373 pairs of isoforms had canonical PAS elements in both isoforms with an average of 246 nt difference between isoforms while 665 pairs had variant PAS elements in both isoforms with an average of 125 nt difference between them. In isoform pairs where the type of PAS element switches, 71% have a shorter isoform with a variant PAS element and a longer isoform with a canonical PAS element with an average of 209 nt between them while the remaining 29% have a canonical PAS element on the shorter isoform and a variant PAS element on the longer isoform with an average of 322 nt between them. B) In genes with three or more 3'-UTR isoforms, genes where the longest and the shortest isoform both have canonical PAS elements have an average of 163 nt between them while genes where the longest and the shortest isoforms both have variant PAS elements have an average of 211 nt between them. 72% of genes switch from a variant PAS elements in the short isoform to a canonical PAS element in the long isoform, with an average of 181 nt between them. 28% of genes have canonical PAS elements in the short isoforms and variant PAS elements in the long 3'-UTR isoform, with an average of 211 nt between the two.

**Supplemental Fig. S6**

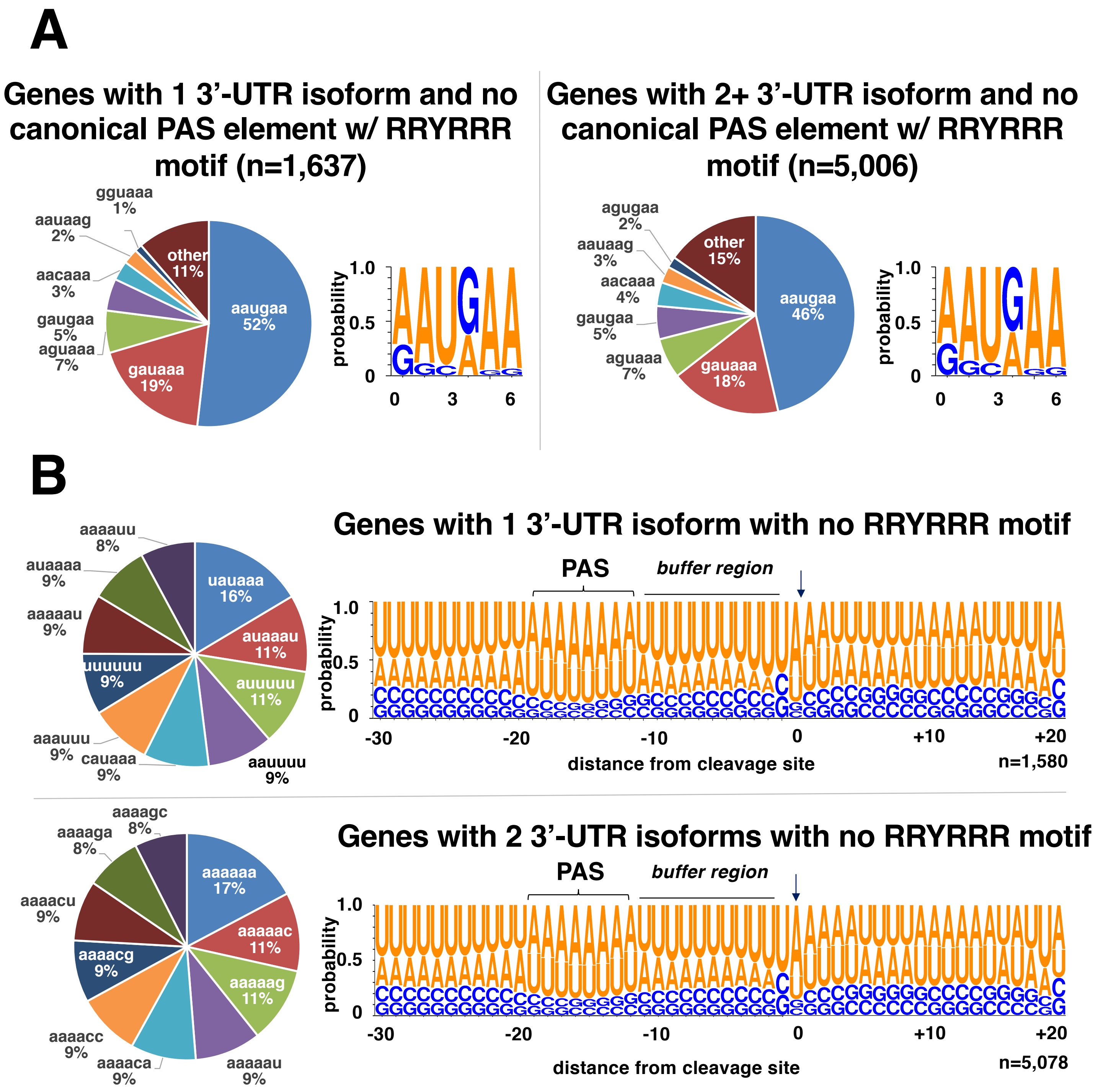


**Supplemental Figure S6: Detection of the PAS element in genes lacking a canonical AAUAAA hexamer.** A) We have searched for the most common RRYRRR motifs located within the last 30 nt in genes with 1 (top) or 2+ (bottom) 3'-UTR isoforms with no canonical PAS and with a detectable RRYRRR motif. The left chart shows the occurrences of the seven most common PAS elements identified in these groups, and the right logo shows the identified PAS motif. Apart from a slight increase in percentage of adenosines and guanosines in 3'-UTRs of genes with 2+ 3'-UTRs, both results are similar. The overwhelming majority of PAS conform with the AAU(G/A)AA with a tolerance of 1 nt purine-purine or pyrimidine-pyrimidine replacement. B) PAS site usage in genes with 1 or 2 3'-UTR isoforms with non canonical PAS and no RRYRRR motif.  The pie chart shows the 10 most common hexamers located within 30 nt upstream of the the cleavage site. Right logo plot shows the nucleotide conservation of the intergenic region encompassing the cleavage site from -30 to +20 nt. Arrow marks the cleavage site. The buffer region and the PAS is marked.

#### Biological Process

##### 1 3'-UTR isoform (n=8,537)

| adj.Pval | nGenes | Pathways |
|---|---|---|
| 5.7e-28 | 165 | phosphorus metabolic process |
| 7.2e-28 | 220 | single-organism metabolic process |
| 2.5e-26 | 159 | phosphate-containing compound metabolic process |
| 8.7e-23 | 149 | macromolecule modification |
| 1.3e-17 | 98 | phosphorylation |
| 1.3e-16 | 130 | cellular protein modification process |
| 1.3e-16 | 130 | protein modification process |
| 4.9e-16 | 164 | cellular protein metabolic process |
| 6.2e-14 | 75 | protein phosphorylation |
| 4.7e-13 | 90 | oxidation-reduction process |

##### 2 3'-UTR isoform (n=4,741)

| adj.Pval | nGenes | Pathways |
|---|---|---|
| 3.6e-20 | 135 | single-organism metabolic process |
| 1.4e-13 | 38 | immune response |
| 1.4e-13 | 38 | immune system process |
| 5.0e-13 | 63 | oxidation-reduction process |
| 8.7e-12 | 35 | innate immune response |
| 7.5e-11 | 82 | phosphorus metabolic process |
| 4.0e-10 | 41 | defense response |
| 1.1e-09 | 78 | phosphate-containing compound metabolic process |
| 9.9e-08 | 41 | carbohydrate derivative metabolic process |
| 1.0e-07 | 51 | small molecule metabolic process |

##### 3+ 3'-UTR isoform (n=1,530)

| adj.Pval | nGenes | Pathways |
|---|---|---|
| 2.3e-03 | 19 | small molecule metabolic process |
| 2.3e-03 | 21 | organonitrogen compound metabolic process |
| 2.3e-03 | 8 | ribonucleoside monophosphate metabolic process |
| 2.3e-03 | 8 | nucleoside monophosphate metabolic process |
| 2.3e-03 | 11 | innate immune response |
| 2.3e-03 | 11 | immune response |

#### Cellular Component

##### 1 3'-UTR isoform (n=8,537)

| adj.Pval | nGenes | Pathways |
|---|---|---|
| 5.4e-28 | 122 | cytoplasmic part |
| 6.1e-22 | 48 | mitochondrion |
| 9.7e-22 | 102 | organelle part |
| 5.2e-18 | 89 | intracellular organelle part |
| 2.5e-15 | 73 | non-membrane-bounded organelle |
| 2.5e-15 | 73 | intracellular non-membrane-bounded organelle |
| 8.7e-13 | 29 | mitochondrial part |
| 1.0e-12 | 94 | nucleus |
| 6.8e-12 | 40 | membrane-enclosed lumen |
| 4.2e-11 | 38 | intracellular organelle lumen |

##### 2 3'-UTR isoform (n=4,741)

| adj.Pval | nGenes | Pathways |
|---|---|---|
| 4.1e-17 | 73 | cytoplasmic part |
| 9.9e-08 | 20 | endoplasmic reticulum |
| 9.9e-08 | 50 | organelle part |
| 4.3e-07 | 21 | mitochondrion |
| 2.9e-06 | 43 | intracellular organelle part |
| 2.9e-06 | 52 | nucleus |
| 3.4e-06 | 23 | extracellular region |
| 4.5e-05 | 23 | endomembrane system |
| 1.9e-04 | 13 | mitochondrial part |
| 1.9e-04 | 7 | membrane raft |

##### 3+ 3'-UTR isoform (n=1,530)

| adj.Pval | nGenes | Pathways |
|---|---|---|
| 4.0e-07 | 25 | organelle part |
| 6.0e-07 | 26 | cytoplasmic part |
| 6.0e-07 | 8 | mitochondrial inner membrane |
| 6.0e-07 | 10 | mitochondrial part |
| 6.0e-07 | 8 | organelle inner membrane |
| 6.0e-07 | 10 | organelle envelope |
| 6.0e-07 | 9 | mitochondrial envelope |
| 6.0e-07 | 10 | envelope |
| 3.5e-06 | 11 | mitochondrion |
| 4.5e-06 | 8 | mitochondrial membrane |

#### Molecular Component

##### 1 3'-UTR isoform (n=8,537)

| adj.Pval | nGenes | Pathways |
|---|---|---|
| 7.6e-64 | 231 | transferase activity |
| 4.2e-54 | 220 | hydrolase activity |
| 4.1e-39 | 204 | nucleotide binding |
| 4.1e-39 | 204 | nucleoside phosphate binding |
| 2.6e-38 | 206 | small molecule binding |
| 6.5e-38 | 195 | anion binding |
| 4.4e-36 | 119 | transferase activity, transferring phosphorus-containing groups |
| 1.1e-29 | 163 | ribonucleotide binding |
| 1.7e-29 | 166 | carbohydrate derivative binding |
| 2.1e-29 | 160 | purine nucleoside binding |

##### 2 3'-UTR isoform (n=4,741)

| adj.Pval | nGenes | Pathways |
|---|---|---|
| 2.4e-28 | 121 | hydrolase activity |
| 1.3e-27 | 117 | transferase activity |
| 1.7e-16 | 56 | oxidoreductase activity |
| 2.2e-15 | 103 | small molecule binding |
| 4.6e-15 | 100 | nucleotide binding |
| 4.6e-15 | 100 | nucleoside phosphate binding |
| 5.3e-15 | 59 | transferase activity, transferring phosphorus-containing groups |
| 1.7e-13 | 70 | zinc ion binding |
| 4.5e-12 | 76 | transition metal ion binding |
| 5.7e-11 | 89 | nucleic acid binding |

##### 3+ 3'-UTR isoform (n=1,530)

| adj.Pval | nGenes | Pathways |
|---|---|---|
| 2.5e-09 | 39 | nucleic acid binding |
| 3.5e-09 | 37 | hydrolase activity |
| 5.3e-08 | 37 | small molecule binding |
| 2.0e-07 | 35 | nucleotide binding |
| 2.0e-07 | 35 | nucleoside phosphate binding |
| 1.1e-06 | 15 | RNA binding |
| 6.0e-06 | 31 | anion binding |
| 1.8e-04 | 26 | carbohydrate derivative binding |
| 2.2e-04 | 21 | zinc ion binding |
| 3.5e-04 | 15 | oxidoreductase activity |

**Supplemental Figure S7: GO term analysis for genes with 1, 2 or 3 3'-UTR isoforms.**

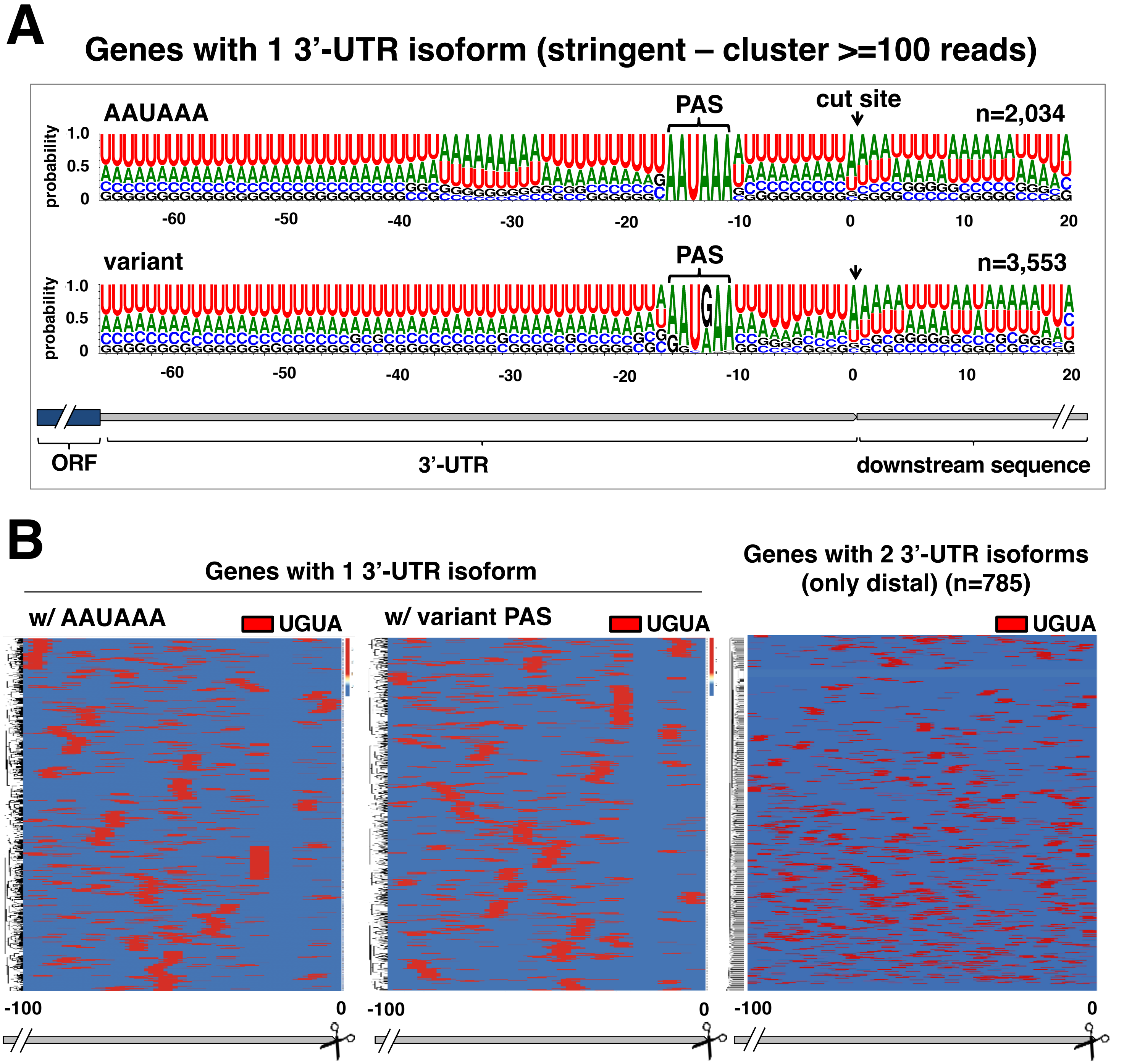


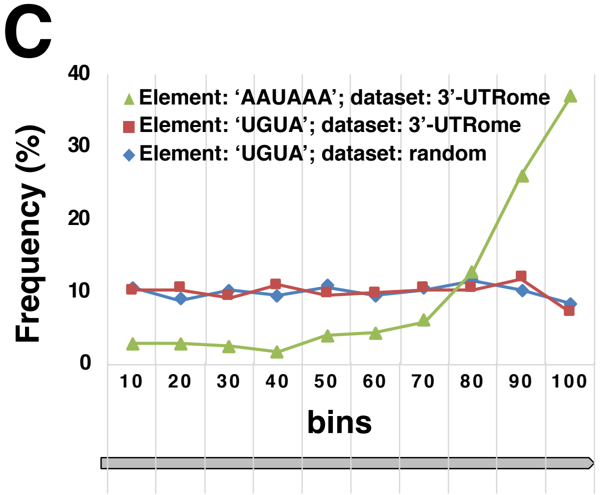


**Supplemental Figure S8: Detection of the 'UGUA' element in *C. elegans* 3'-UTRs.** A) Logo plot of the transcript's region within the cleavage site in genes with only one 3'-UTR isoform and with a canonical or variant PAS element. B) Identification of the 'UGUA" motif (red) within 100 nt upstream of the cleavage site in genes with one or two 3'-UTR isoforms. C) Binned frequency distribution of the occurrences of the AAUAAA and UGUA elements in distal 3'-UTR isoforms as in the right heatmap in Panel B (green and red) vs. the occurrences of the UGUA motif in a randomly generated 3'-UTR dataset (blue) (n=785).

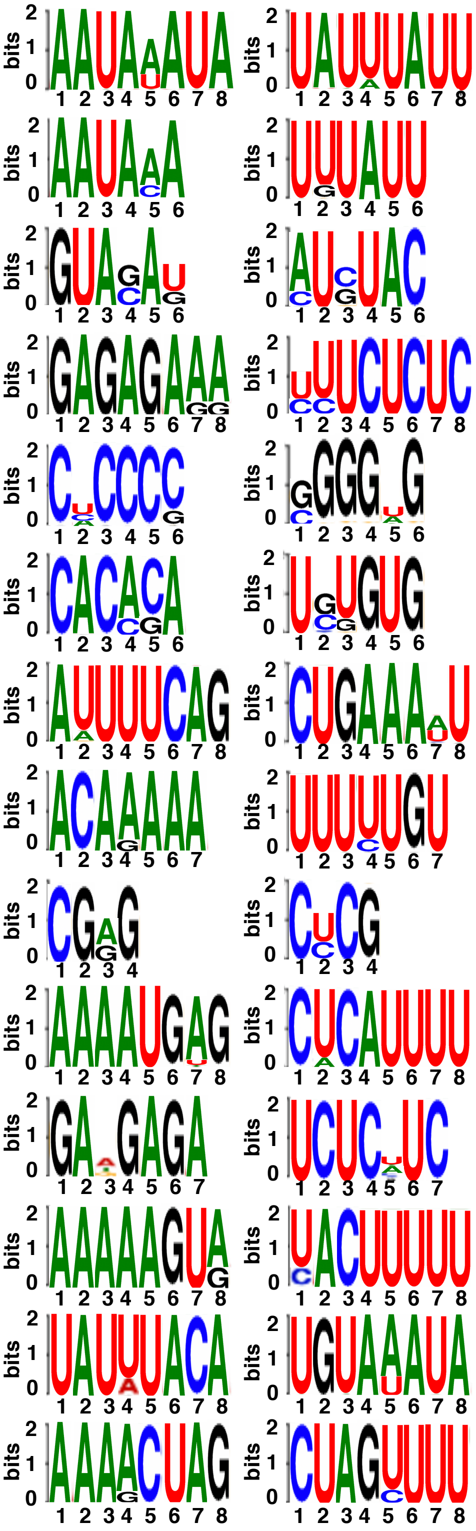

| | Motif | Logo | RC Logo | E-value | Unerased E-value |
|---|---|---|---|---|---|
| 1. | AATAWATA | | | 4.6e-027 | 4.6e-027 |
| 2. | AATAMA | | | 1.2e-010 | 8.8e-015 |
| 3. | GTASAK | | | 1.2e-009 | 8.6e-010 |
| 4. | GAGAGARR | | | 4.7e-008 | 6.6e-008 |
| 5. | CHCCCS | | | 2.2e-007 | 1.8e-007 |
| 6. | CACMSA | | | 3.8e-006 | 1.7e-007 |
| 7. | ADTTTCAG | | | 4.8e-005 | 7.6e-005 |
| 8. | ACARAAA | | | 1.4e-004 | 7.2e-007 |
| 9. | CGRG | | | 1.6e-003 | 2.7e-006 |
| 10. | AAAATGDG | | | 6.8e-003 | 4.0e-004 |
| 11. | GADGAGA | | | 4.4e-003 | 2.5e-004 |
| 12. | AAAAAGTR | | | 1.7e-002 | 1.9e-003 |
| 13. | TATWTACA | | | 1.9e-002 | 9.8e-004 |
| 14. | AAARCTAG | | | 2.6e-002 | 4.5e-002 |

**Supplemental Figure S9: Detection of enriched elements in *C. elegans* 3'-UTRs of genes with 2 3'-UTR isoforms (only distal) (n=785).** These motifs have been detected using the meme suite (Bailey et al., 2015.)

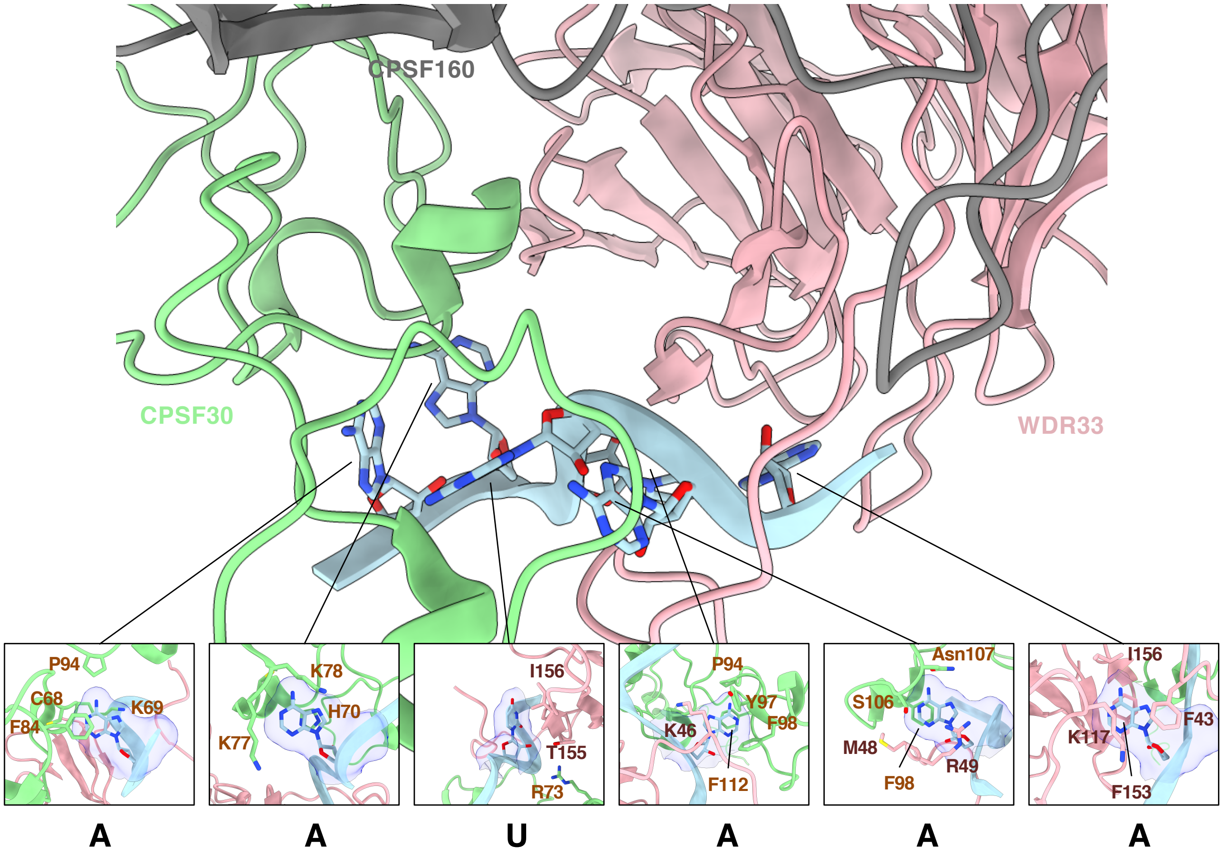


**Supplemental Figure S10: Nucleotide binding site of the human CPSF160-WDR33-CPSF30 complex.** Ribbon representation of the cryo-EM structure of human CPSF160-WDR33-CPSF30 complex (PDB code: 6DNF) (Sun et al., 2018). The nucleotides of the bound RNA fragment do not show a specific interaction with either CPSF30 or WDR33. The interactions are mostly established by π-π ring stacking. Color gray shows the CPSF160, pink for WDR33, and light green for CPSF30. Sticks represent the RNA molecules bound with CPSF30 and WDR33. Surfaces in the inlets are for individual nucleotides.

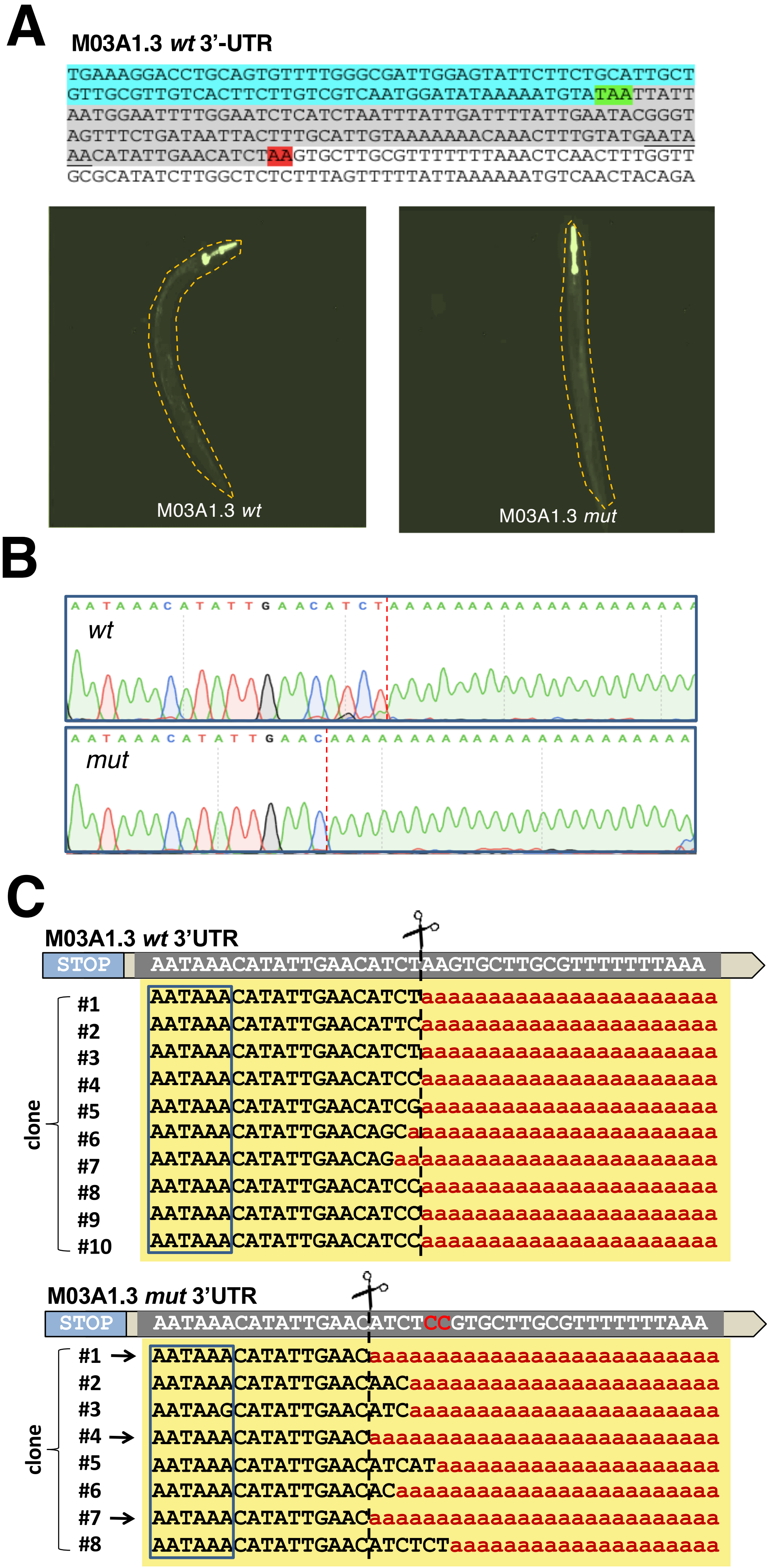


**Supplemental Figure S11: *In vivo* cleavage assay for M03A1.3.**

A) M03A1.3 genomic region cloned downstream of the GFP reporter. Blue: terminal portion of the M03A1.3 ORF. Green: STOP codon. Gray: 3'-UTR. Red: mutated terminal adenosine nucleotides. The transgenic worms expressing the Pmyo-3::GFP::M03A1.3_3'-UTR *wt* and mutant cassette are shown below.

B) At the completion of the experiment, we recovered the total RNA and performed RT-PCR experiments using a forward primer annealing within the GFP ORF and a reverse polydT primer with two anchors containing Invitrogen Gateway adapters. The resultant amplicons were then subcloned in gateway vectors and sequenced to detect the cleavage site. An example of resultant trace files is shown.

C) Examples of 10 clones identified in this study for M03A1.3. The removal of the terminal genomic adenosine nucleotide induces a cleavage site 3 nt upstream of the canonical cleavage site in three clones (arrows), which also contain a terminal adenosine nucleotide. The PAS element is boxed in blue color.

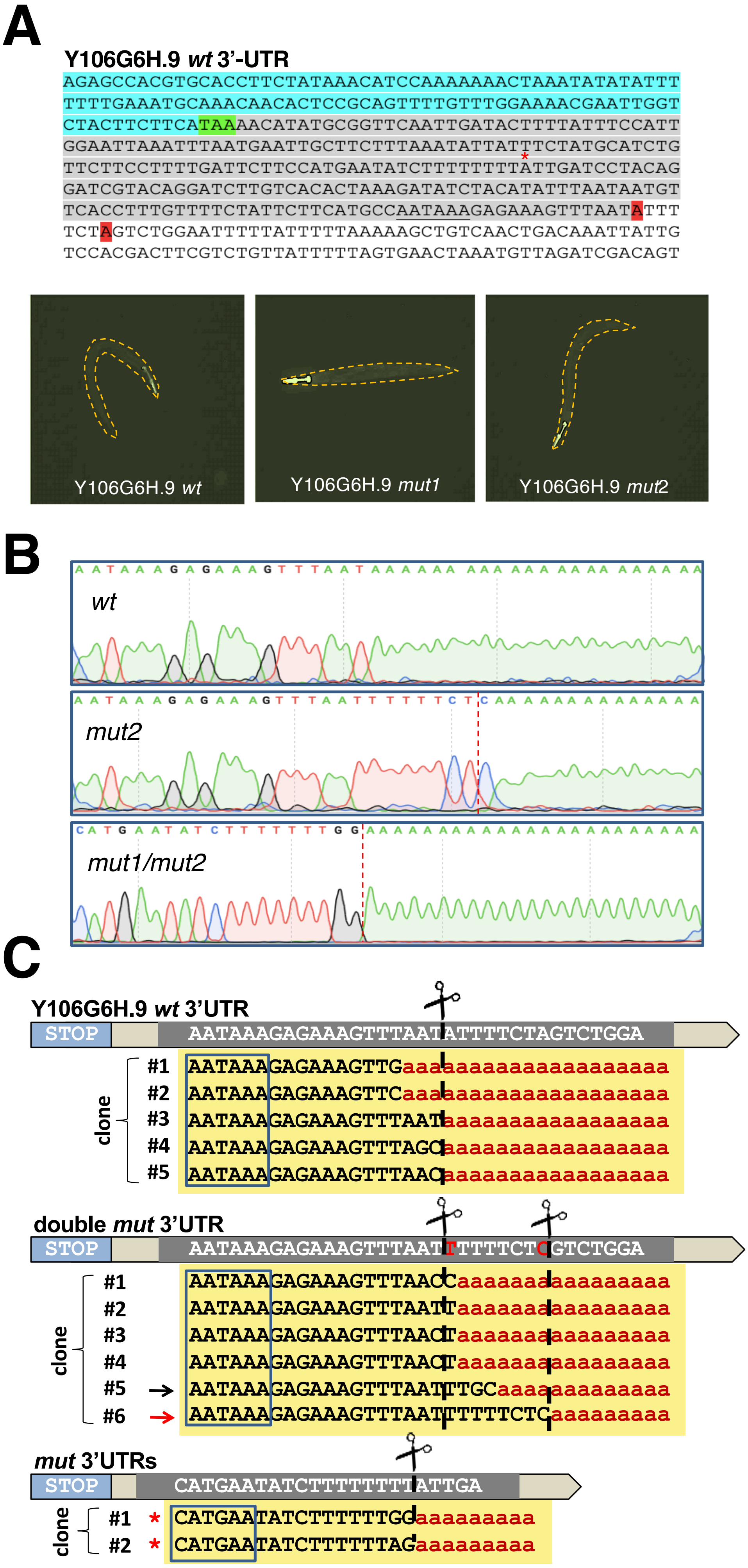


**Supplemental Figure S12: *In vivo* cleavage assay for Y106G6H.9.**
A) Y106G6H.9 genomic region cloned downstream of the GFP reporter. Blue: terminal portion of the Y106G6H.9 ORF. Green: STOP codon. Gray: 3'-UTR. Red: mutated terminal adenosine nucleotides. Red Asterisk: position of the cryptic cleavage site (see below). The transgenic worms expressing the Pmyo-3::GFP:: Y106G6H.9_3'-UTR *wt* and mutant cassette are shown below.
B) At the completion of the experiment we recovered the total RNA and performed RT-PCR experiments using a forward primer annealing within the GFP ORF and a reverse polydT primer with two anchors containing Invitrogen Gateway adapters. The resultant amplicons were then subcloned in gateway vectors and sequenced to detect the cleavage site. An example of resultant trace files is shown.
C) Examples of several clones identified in this study for Y106G6H.9. In the *wt* we were able to detect two classes of cleavage sites, both ending within 4 nt of each other with a terminal adenosine nucleotide. In the double mutant, the removal of the terminal genomic adenosine induces a cleavage skip in two clones (arrows). In one case (red arrow), the cleavage occurs 20 nt downstream of the PAS element. Two of the mutant clones also shown an occurrence of a new cryptic cleavage site 100 nt upstream of the natural site (red asterisks), which also contain a terminal adenosine nucleotide at their 3'end. The PAS element is boxed in blue color.

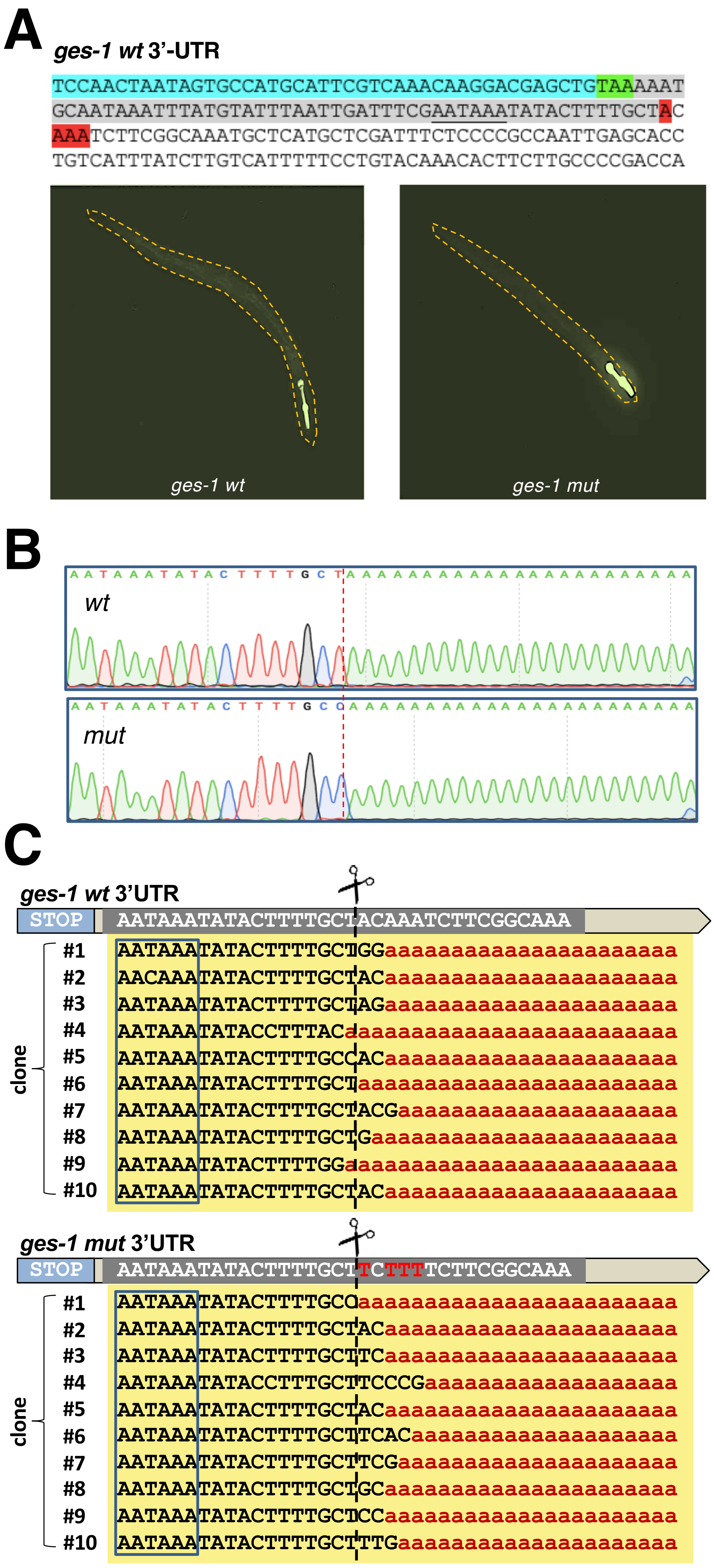


**Supplemental Figure S13:** ***In vivo* cleavage assay for *ges-1*.**
A) *ges-1* genomic region cloned downstream of the GFP reporter. Blue: terminal portion of the *ges-1* ORF. Green: STOP codon. Gray: 3'UTR. Red: mutated terminal adenosine nucleotides. The transgenic worms expressing the Pmyo-3::GFP::ges-1_3'-UTR *wt* and mutant cassette are shown below.
B) At the completion of the experiment we recovered the total RNA and performed RT-PCR experiments using a forward primer annealing within the GFP ORF and a reverse polydT primer with two anchors containing Invitrogen Gateway adapters. The resultant amplicons were then subcloned in gateway vectors and sequenced to detect the cleavage site. An example of resultant trace files is shown.
C) Examples of 10 clones identified in this study for *ges-1*. The removal of the terminal genomic adenosine nucleotide does not alter the cleavage site but makes it more variable. The PAS element is boxed in blue color.

**A**

**GO Term: Biological Process**

| adj.Pval | nGenes | Pathways |
|---|---|---|
| 7.2e-05 | 7 | oxidation-reduction process |
| 5.9e-04 | 2 | deoxyribonucleoside diphosphate metabolic process |
| 1.7e-03 | 8 | single-organism metabolic process |

**GO Term: Molecular Component**

| adj.Pval | nGenes | Pathways |
|---|---|---|
| 1.2e-03 | 5 | oxidoreductase activity |

**Functional Enrichment: Kegg**

| adj.Pval | nGenes | Pathways |
|---|---|---|
| 1.7e-03 | 1 | Pentose and glucuronate interconversions |
| 1.7e-03 | 1 | Fructose and mannose metabolism |

**B**

| miRNA | Sequence | Hits |
|---|---|---|
| miR-272 | uguaggcaugggguuug | 7 |
| miR-2217a | cagaguggggcagucggugucgauc | 6 |
| miR-2217b | cagagcggggcagucggugucaauc | 6 |
| miR-5553 | ucaaugggguagcacguggcaaga | 6 |
| miR-265 | ugagggaggaagggugguau | 5 |
| miR-34 | aggcagugugguuagcugguug | 5 |
| miR-44 | ugacuagagacacauucagcu | 5 |
| miR-4935 | gccggcgagagagguggagagcg | 5 |
| miR-795 | ugagguagauugaucagcgagcuu | 5 |
| miR-8190 | cgggaaaucgcuuuggaauccagga | 5 |
| miR-8194 | augcgccuuuaaaaagguacgg | 5 |
| miR-1822 | aguuucucugggaaagcuaucggc | 4 |
| miR-71 | ugaaagacaugggguagugagacg | 4 |

**Supplemental Figure S14: miRNA target analysis in genes with 2 3'-UTR isoforms which either gain or lose a miRNA binding site.** A) GO Term analysis of genes with multiple 3'-UTRs which gain or lose a miRNA target as predicted using our miRanda 'stringent' dataset (n=132). B) Most common miRNAs, their sequences, and number of occurrences.

#### Additional Materials and Methods

##### Comparative analysis of *C. elegans* members of the CPC

We have downloaded the protein sequences of each known member of the human CPC and used BLAT algorithm to identify *C. elegans* genes with high homology to their human counterparts. We then performed a protein BLAST analysis using the tools available at the NCBI website to obtain the amino acid sequences for the fly, rat, and mouse orthologs. These amino acid sequences were then aligned using Clustal Omega Multiple Sequence Alignment with standard parameters. At the completion of the analysis, we used the Batch NCBI Conserved Domain Search (Batch CD-Search) against the database CDD- 52910 PSSMs using standard parameters to identify the conserved domains across the aligned protein sequences. We then used these results to populate the location of these elements within the alignment shown in **Supplemental Figure S1**.

##### Plasmid DNA isolation, sequencing and visualization

All plasmids used in this study were prepared from cultures grown overnight in LB using the Wizard Plus SV Minipreps DNA Purification System (Promega) according to the manufacturer’s instructions. DNA samples were sequenced with Sanger sequencing performed at the DNASU Sequencing Core Facility (The Biodesign Institute, ASU, Tempe, AZ).

**RNAi experiments**

RNAi experiments were performed in standard NGM agar containing 1mM IPTG and 50 μg/ml ampicillin. These plates were seeded with 75 μl of RNAi clone bacteria and allowed to induce for a minimum of 16 hours. 5 N2 *C. elegans* at the L1 stage were aliquoted for each RNAi clone tested. Three days after plating, the progeny was scored for embryonic lethality. Each RNAi experiment was performed in triplicate. The total number of hatched and not hatched eggs was the following: *cpsf-1(CPSF160)* n=567*; cpsf-2(CPSF100)* n=557*; cpsf-4(CPSF30)* n=1,251*; cpf-2(CstF64)* n= 716*; cpf-1(CstF50)* n=801*; cfim-1(CFIm25)* n=644*; cfim-2(CFIm68)* n=739*; lrp-2(CFIm59)* n=1,120*; symk-1(symplekin)* n=208*; tag-214(RBBP6)* n=753*; pcf-11(CPF11)* n=428*; clpf-1(CLP1*) n=841*.*

**Mutagenesis of 3'-UTRs cleavage sites**

The mutagenesis reactions to remove the adenosine nucleotides near the cleavage sites were carried out using the QuikChange Site-Directed Mutagenesis Kit (Agilent). The mutagenesis DNA primers for the site mutation reactions are available in **Supplemental Table S2**. Each mutagenesis reaction was followed by DNA digestion using Dpn-1 enzyme and transformed in Top10 competent cells (Thermo Fisher Scientific) in agar plates containing 20mg/μL of kanamycin. We validated the nucleotide mutation using Sanger sequencing approach. *Wild type* and mutant 3'-UTRs cloned in pDONR P2RP3 were then shuttled into destination vectors using the Gateway LR Clonase II Plus Enzyme Mix (Invitrogen, Carlsbad, CA). The finalized destination vectors contained the *C. elegans* pharynx promoter (Pmyo-2) in the first position, a GFP sequence with a mutated STOP codon in the second position, and the *wt* or mutant 3'-UTRs used in this study in the third position. The resultant recombined constructs were then transformed in Top10 competent cells (Thermo Fisher Scientific) and plated on 10mg/μL ampicillin plates overnight. The success of the recombination reaction was confirmed using Sanger sequencing with the M13F DNA primer.

**Preparation of transgenic worm lines**

EG6699 strain worms were kindly provided by Christian Frokjaer-Jensen (Frokjaer-Jensen et al. 2008). These worm strains were maintained at 18°C on nematode growth media (NGM) agar plates and propagated on plates seeded with OP50-1 bacteria. To synchronize worms for injections, EG6699 worms were bleached with bleaching solution (1 M NaOH) four days before injections. Each construct was mixed with an injection master mix containing pCFJ601 (25 ng/μl), pgH8 (10 ng/μl), and pCFJ104 (5 ng/μl) vectors. Injection needles were loaded with the injection mixture and mounted to the Leica DMI300B microscope. The needle was pressurized with 22 psi through the FemtoJet (Eppendorf). Young adult EG6699 worms were picked onto an agarose pad covered with mineral oil on a glass coverslip. Injected worms were rescued onto an NGM plate and rinsed with M9 buffer. Two days post-injections, the F1 progeny were screened with a Leica DMI3000B microscope for both *unc-119* rescues and expression of the red fluorescence produced by the co-injection marker and then isolated onto individual plates. These worms were allowed to lay eggs, and then the F2 progeny was screened for fluorescence. Once 75% of the progeny on a single plate were transgenic, the strains were used for further experimentation.

**Worm genotype validation**

Populations obtained from single worms from each of the seven strains were lysed using worm lysis buffer (EDTA, 0.1 M Tris, 10% Triton-X, Proteinase K, 20% Tween 20). These worms were subjected to heating in a Bio-Rad T100 Thermal Cycler. To confirm that the mutated cleavage site was present in the injected strains, we used PCR approach using Platinum Taq polymerase (Invitrogen) with a forward DNA primer binding the beginning of the GFP sequence and 3'-UTR-specific reverse DNA primers. The PCR product was then sequenced using Sanger sequencing with a forward DNA primer binding to the GFP sequence present in the injected construct.

**Detection of the 3'-UTR cleavage skipping**

Total RNA was extracted from transgenic strains using the Direct-zol RNA MiniPrep Plus kit (RPI) according to the manufacturer's instructions. We tested approximately 10 independent *wt* and mutant clones for each 3'-UTR. Approximately 50 μL of worm pellet was used for extraction. cDNA was synthesized using a reverse transcription reaction using Superscript II enzyme (Invitrogen). The first strand reaction was performed using a reverse poly dT DNA primer containing two anchors and the attB Gateway BP recombination element (Invitrogen). The second strand of the cDNA was synthesized using a PCR with HiFi taq polymerase (Thermo Fisher Scientific) and the forward DNA primer containing the pDONR P2RP3 Gateway element (Invitrogen), which binds to GFP and the same reverse poly dT DNA primer used in the first strand reaction. The BP Gateway kit (Invitrogen) was once again used to clone the cDNA which contains the polyA tail into pDONR P2RP3. These constructs were then transfected into Top10 competent cells (Thermo Fisher Scientific) and plated on agar plates containing 20mg/μL of Kanamycin. About 8-10 colonies were then sequenced with Sanger sequencing using the M13F DNA primer to map the location of the cleavage site.

**Updated miRanda Predictions**

We downloaded a complete list of *C. elegans* miRNAs from miRBase (Griffiths-Jones et al. 2006) and the miRanda algorithm v3.3a (John et al. 2004) from the microrna.org website. We queried the 3'-UTRome v2 with the miRanda algorithm using both standard and stringent parameters. The stringent query used was '-strict -sc -1.2'. The standard query produced 58,330 putative miRNA targets; the stringent query produced 12,136 putative miRNA targets. Both these predictions are included in WormBase (Lee et al. 2018) as individual tracks.

**Homology model building**

Homology modeling was performed using SWISS_MODEL (Waterhouse et al. 2018) with a matched templated of human CPSF160-WDR33-CPSF30 complex (PDB code: 6DNF) (Sun et al. 2018). The molecular graphics were prepared using the UCSF ChimeraX software (version 0.8) (Goddard et al. 2018).
